# Subcortical origin of nonlinear sound encoding in auditory cortex

**DOI:** 10.1101/2024.02.01.578402

**Authors:** Michael Lohse, Andrew J King, Ben DB Willmore

## Abstract

A major challenge in neuroscience is to understand how neural representations of sensory information are transformed by the network of ascending and descending connections in each sensory system. By recording from neurons at several levels of the auditory pathway, we show that much of the nonlinear encoding of complex sounds in auditory cortex can be explained by transformations in the midbrain and thalamus. Modeling cortical neurons in terms of their inputs across these subcortical populations enables their responses to be predicted with unprecedented accuracy. In contrast, subcortical responses cannot be predicted from descending cortical inputs, indicating that ascending transformations are irreversible, resulting in increasingly lossy, higher-order representations across the auditory pathway. Rather, auditory cortex selectively modulates the nonlinear aspects of thalamic auditory responses and the functional connectivity between subcortical neurons, without affecting the linear encoding of sound. These findings reveal the fundamental role of subcortical transformations in shaping cortical responses.

## Introduction

Sensory systems need to represent the external environment in ways that allow animals to use this information for survival. The dominant tools for characterizing the tuning properties of individual sensory neurons are spatiotemporal (visual) and spectrotemporal (auditory) receptive fields (STRFs). These models provide a linear mapping from the spatio/spectrotemporal variables in the environment to neuronal activity and are reasonably successful at describing and predicting neuronal responses at lower levels of the visual and auditory pathways. For example, measuring STRFs provides an effective way of characterizing the tuning properties of auditory nerve fibers^1^.

At higher levels of the sensory pathways, however, neural representations are seemingly more abstract, and correspondingly harder to model^2,3^. It is well established that STRF models perform relatively poorly in sensory cortex, most likely because cortical coding takes place in complex networks of highly interconnected excitatory and inhibitory cells^4–6^. However, it has proven challenging to produce nonlinear models that capture cortical sensory processing as accurately as STRF models capture the behaviour of subcortical neurons. Moreover, and particularly in the auditory system, it is not known to what degree the coding of complex stimuli in the cortex reflects subcortical transformations of the input.

To better understand how auditory neurons represent the environment, much effort has been made to expand STRF models – for example, by incorporating sensory and behavioural contexts^7–14^, as well as by exploring different nonlinearities^9,15–18^. While these approaches have provided insights into what features sensory neurons may represent, they have yet to result in a generally accepted model of how cortical auditory representations differ from subcortical representations or how and where neuronal representations of sounds in the auditory pathway are nonlinearly transformed into the complex representations found in cortex. This is partly because the shapes of the STRFs along the ascending auditory pathway are remarkably similar, in contrast to the qualitative transformations that take place in the visual system^19–21^.

However, it also results from the difficulty of fitting ever more complex models to limited physiological datasets. As the complexity of the models increases, the ability of a limited dataset to constrain these models precisely decreases, resulting in suboptimal predictions. As a result, models of auditory cortical neurons fail to capture much of their stimulus-locked variability. It is not clear to what degree this arises from nonlinear processing, variability in the responses or imprecision in cortical encoding.

The cortex returns descending projections to nearly all subcortical levels, including the thalamus and midbrain. The excitability and response properties of neurons at different subcortical levels can be altered by manipulating the activity of neurons in the auditory cortex (reviewed in ^22–24^), and there is growing evidence that these descending corticofugal projections play important roles in auditory perception^25,26^ and learning^27^. However, it is not yet known how these projections contribute to linear or nonlinear subcortical sensory encoding.

Here, we investigate the origin of nonlinear representations of sound in the auditory cortex. By exploring the nature of ascending and descending population communication between the auditory midbrain, thalamus and cortex, we show that non-linear subcortical transformations profoundly shape the response properties of cortical neurons. By combining this approach with optogenetic manipulation of cortex, we further demonstrate that cortex selectively modulates the nonlinear representations in thalamus and the functional connectivity between subcortical neurons, without affecting the linear encoding of sound. Furthermore, this new population communication model allows us to predict to an unprecedented degree the sound-evoked responses of cortical neurons.

## Results

To understand how encoding of sound in auditory cortex arises as a function of inputs from populations of other auditory neurons, we recorded extracellular responses from 1403 units (from 31 mice) to identical complex sounds at three levels of the auditory system: the midbrain (inferior colliculus, IC, *n* = 432), thalamus (medial geniculate body, MGB, *n* = 355), and primary auditory cortex (A1, *n* = 616). We developed a population communication model (PCM), which describes the sound-locked activity of individual units in terms of the sound-locked activity of populations of other units in the auditory system (**Figure 1A**). The population communication model is a generalized linear-nonlinear encoding model, involving regularized linear regression between the response of the target unit and a set of input signals at multiple time lags, followed by a static nonlinearity (see *Methods*). The input signals are the recorded responses of other units, enabling us to directly probe the transformations of information that occur between the input units and the target unit.

**Figure 1:**
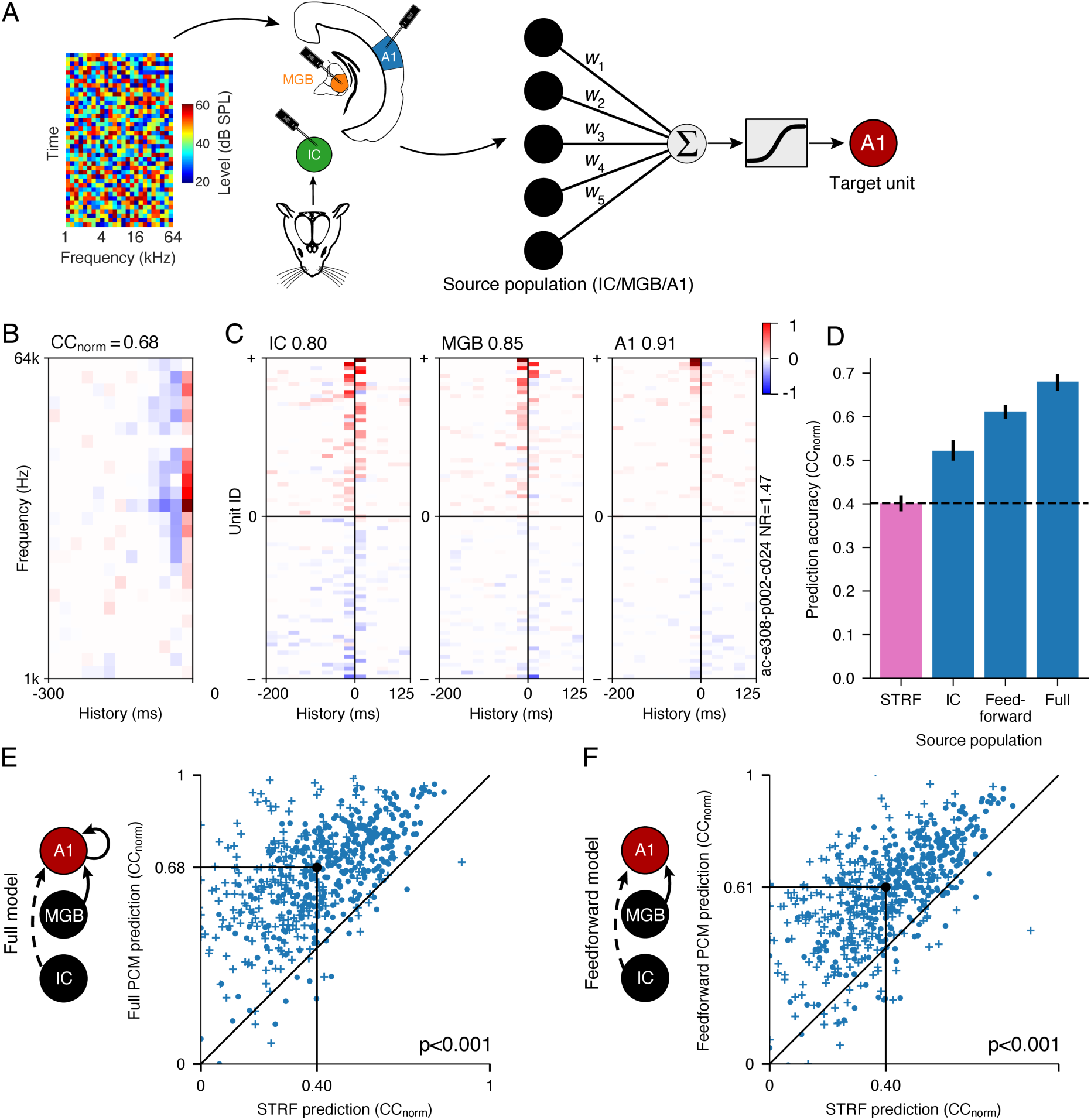
Feedforward population communication greatly improves the predictability of auditory cortical responses compared to receptive field models. (A) Schematic showing brain areas recorded from in response to dynamic random chords and the population communication model (PCM) used to explore communication between populations of neurons. Neurons in one or more areas (the source population) are used as input to the model, which is trained to describe the responses of each neuron (target unit) in the dataset. (B) Example of an STRF from a unit in A1. (C) Examples of PCM weights that describe the responses of the same A1 unit as in B. Each set of PCM weights describes an optimised linear mapping between the responses of the source population and the responses of the target unit. (D) Population communication models (blue bars) greatly outperform STRF models (pink bar), and the predictability of auditory cortical increases as more nonlinear transformations across the ascending pathway are included (*p* < 0.001; *n* = 616 units). Error bars are 95% nonparametric confidence intervals. (E) Comparison of the prediction performance of auditory cortical units between the full PCM (i.e., using populations from IC, MGB and A1 to capture the variability of auditory cortical neurons) and the STRF (*p* < 0.001; *n* = 616 units). Black dot indicates medians. (F) as in E, but comparing feedforward PCM predictions (i.e., using populations from IC and MGB to capture the variability of auditory cortical neurons). Plusses denote units recorded under anaesthesia, filled circles denote units recorded in awake animals.

### Subcortical transformations account for nonlinear auditory cortical responses

We first asked whether the population communication model could accurately capture the responses of auditory neurons at higher levels of the auditory pathway. To do this, we trained the model to describe the responses of each recorded unit in A1 in terms of the mean responses of all non-simultaneously recorded units (enabling us to focus on sound-locked activity, excluding any contribution from noise correlations between simultaneously recorded units). We also trained a standard linear-nonlinear STRF model (**Figure 1B**), and a nonlinear neural network receptive field model (NRF model^18^) on the same units for comparison. We compared the ability of the three models to predict real neuronal responses by measuring the normalized correlation coefficient (CC_norm_^28^) on a held-out dataset.

We found that a population communication model predicting cortical neural activity from populations of units in IC, MGB and A1 (‘Full’ model) substantially outperformed both the STRF model and nonlinear NRF model. Specifically, the population communication model captures 69.4% (CC_norm_^IC+MGB+A1^/CC_norm_^STRF^: 0.68/0.40) and 76.4% (CC_norm_^IC+MGB+A1^/CC_norm_^NRF^: 0.68/0.38) more of the variance of auditory cortical responses, compared to STRF and NRF models, respectively (**Figures 1C-E and S1;** *p* < 0.001, *n =* 616). This both provides a new lower bound on the proportion of the variance of A1 responses that can be captured by models and quantifies the balance between the linearity and nonlinearity of A1 neurons for the first time (**Figures 1E and S2**). Importantly, it demonstrates that a major part of auditory cortical responses to complex sounds is of a nonlinear nature that cannot be captured by STRFs or nonlinear neural network models.

The high performance of the full population communication model (i.e., predicting using IC+MGB+A1 units) could result from different cortical units having similar responses, or because the responses of cortical neurons are well described by patterns of feedforward connectivity from subcortical neurons. We therefore asked to what degree a purely feedforward model (with inputs from IC and MGB) could account for this drastic improvement in our ability to predict A1 neural activity. We found that, as expected, the full model (**Figure 1E)** significantly outperformed the purely feedforward model (**Figure 1F**, *p* << 0.001, *n =* 616), demonstrating spectrotemporal nonlinearity introduced by A1 itself. Surprisingly, however, the difference in performance was fairly small, indicating that most of the superiority of the full population communication model over the STRF can be achieved using only subcortical feedforward inputs to the auditory cortex (**Figure 1F**). This demonstrates that a large proportion of the nonlinearity in cortical responses (i.e., the variance unaccounted for by STRF models) actually arises because of nonlinear feedforward transformations in the subcortical ascending auditory pathway.

### Cortical responses arise from successive nonlinear transformations

We then set out to investigate where these nonlinear transformations occur along the subcortical ascending auditory pathway, and how information is passed on to higher levels of auditory processing. As expected, we found that the predictability of auditory responses to complex sounds by STRFs decreased with every step along the auditory pathway (**Figures 2A,B and S3**, *p* < 0.001, *n*_IC_ *=* 432*, n*_MGB_ *=* 355*, n*_A1_ *=* 616), suggesting a progressive decrease in the linearity of spectrotemporal representation. This is not surprising since the IC is generally considered to be a processing stage with a more faithful representation of the spectrotemporal content than A1^29,30^. What is unexpected, however, is that A1 responses are better predicted by the full population communication model than IC responses are by the STRF model, suggesting that A1 has a much more faithful - albeit nonlinear - representation of the spectrotemporal content of the sound than previously appreciated (**Figures 1D, 2B, and S4**).

**Figure 2:**
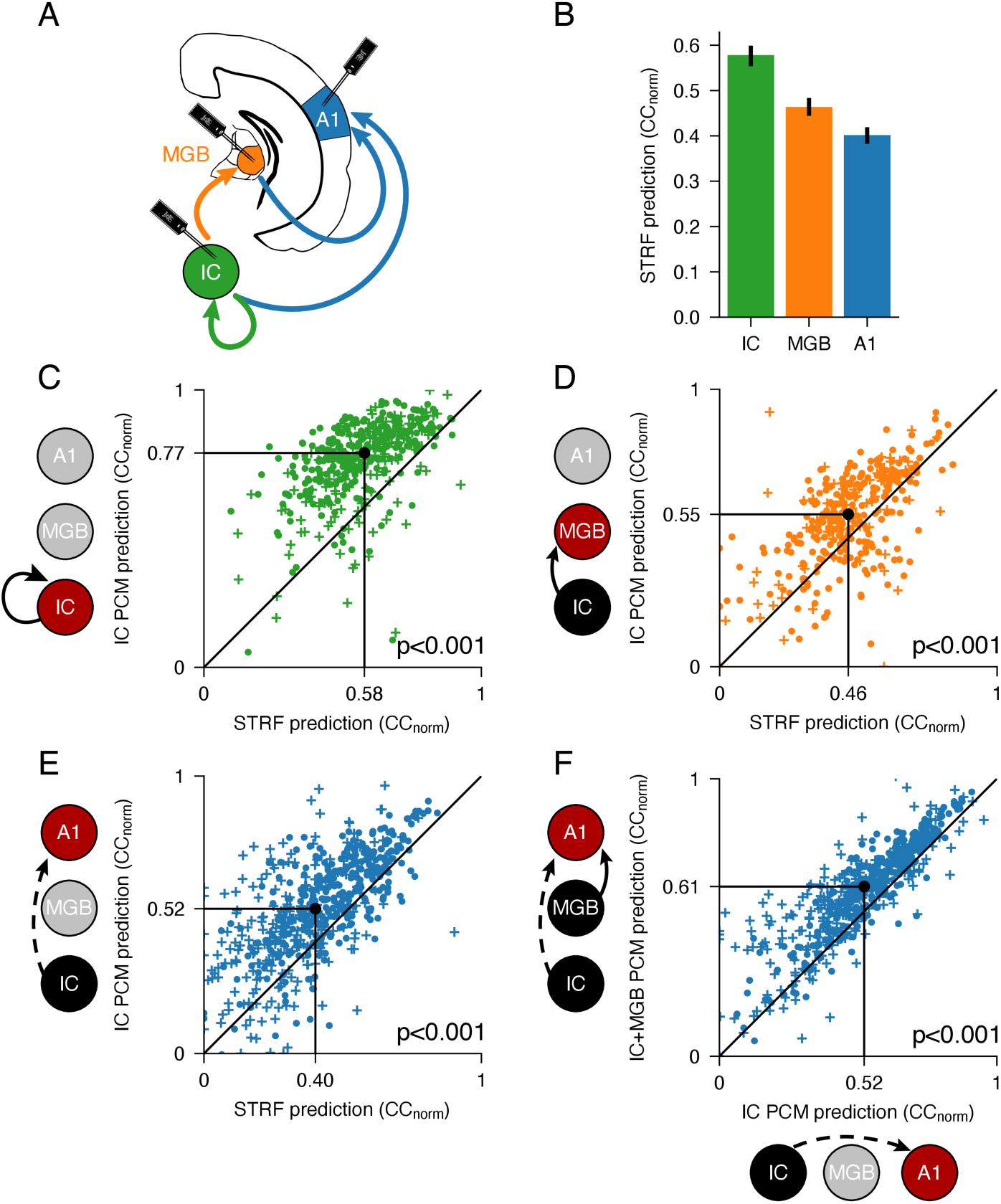
Representations of sound are nonlinearly transformed at each stage of the auditory pathway. (A) Schematic of recording sites in the IC, MGB and A1. Arrows indicate processing levels between which population communication models were applied. (B) Decreasing capacity of STRF models to predict neuronal responses across the ascending auditory pathway. Error bars are 95% nonparametric confidence intervals. (C) Comparison of prediction accuracy of IC responses between the population communication model (using IC units as the source population) and STRF models of the same units. Black dot indicates medians. (D) as in C, but predicting MGB unit activity (using IC as the source population). (E) as in C, but predicting A1 unit activity (using IC as the source population). (F) Prediction accuracy of A1 responses for the population communication model using both IC and MGB units as the source population (feedforward model) and the population communication model using IC units only as the source population. Plusses denote units recorded under anaesthesia, filled circles denote units recorded in awake animals.

Strikingly, we also found that predictions of IC unit activity by local IC population communication markedly outperformed STRF predictions of IC units (**Figure 2C**, *p* < 0.001, *n =* 432), indicating that a major nonlinear transformation of spectrotemporal information has already taken place by the time auditory signals are represented at the level of the midbrain. These midbrain spectrotemporal representations are then passed on via the thalamus to cortical neurons through population communication, as demonstrated by the capacity of population communication from midbrain units to significantly improve predictions of both thalamic (**Figure 2D**, *p* < 0.001, *n =* 355) and cortical (**Figure 2E**, *p* < 0.001, *n =* 616) responses compared to STRF models.

Importantly, adding MGB units to the IC-only model (i.e., the feedforward model) further increased our ability to predict auditory cortical responses (**Figure 2F**, *p* < 0.001, *n =* 616). This suggests that an additional nonlinear transformation of acoustic information takes place in the thalamus, and that this, in turn, contributes to the representation of complex sounds in A1.

Together, these findings demonstrate that nonlinear transformations take place at multiple steps along the ascending auditory pathway. The results of these transformations are passed on from level to level through population communication between hierarchical levels of the auditory pathway, ultimately shaping cortical representations of spectrotemporal information.

### Cortical representations of sound are higher order and lossy

We next assessed the role of descending population communication in shaping subcortical auditory spectrotemporal representations, using models based on the responses of higher-level units as input for modelling lower-level units. We found that a model using cortical units as input (A1 PCM) poorly predicted both MGB and IC responses (**Figure 3A-C**), with STRF models of IC responses substantially outperforming the predictions of the cortical population communication model (**Figure 3B**, *p* < 0.001, *n =* 432). Indeed, we found that subcortical representations could not be captured well by descending cortical population communication, and that the more ascending steps away the predicting population is, the worse the prediction becomes (**Figures 3C,D and S5**, IC vs MGB, MGB vs A1, IC vs A1: all p << 0.001, *n*_IC_ *=* 432*, n*_MGB_ *=* 355*, n*_A1_ *=* 616), even when the longer latencies of cortical responses are taken into account (**Figure S6**). This suggests that the nonlinear transformations taking place along the ascending auditory pathway are irreversible: a higher level (e.g., cortex) representation cannot be transformed into a lower level (e.g., midbrain) representation. This implies that cortical representations are higher order, with information being discarded along the ascending auditory pathway.

**Figure 3:**
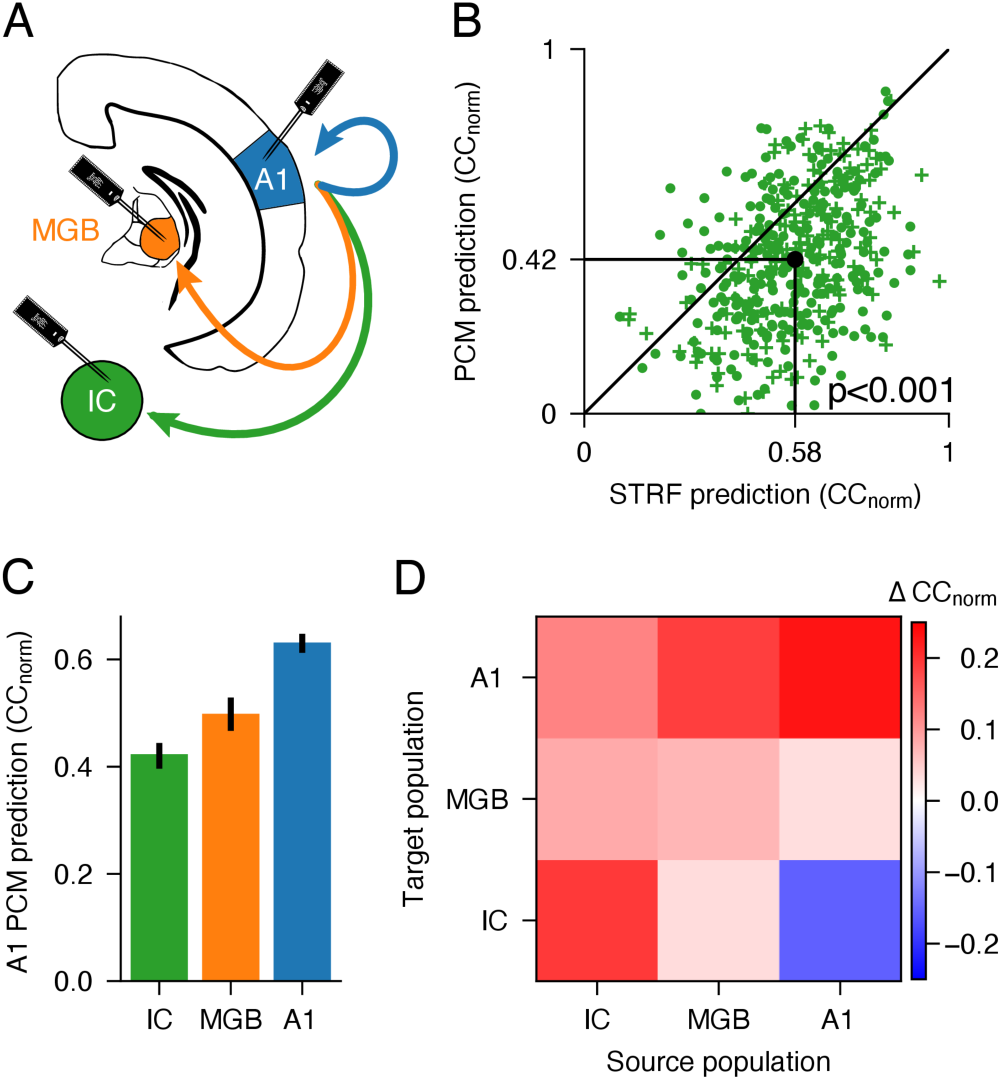
Nonlinear transformations along the ascending pathway are irreversible. (A) Schematic of recording sites and descending connections modelled. (B) Comparison of prediction accuracy of IC unit responses for a population communication model in which A1 units were used as inputs versus STRF models of the same IC units. Black dot indicates medians. Plusses denote units recorded under anaesthesia, filled circles denote units recorded in awake animals. (C) Decreasing capacity of A1 units to predict neuronal responses at progressively earlier stages in the auditory pathway. Error bars are 95% nonparametric confidence intervals. (D) Difference (median) between prediction accuracy of population communication models and STRF models for units recorded in the IC, MGB and A1 (target population) with regressor inputs (source population) from different processing levels (see also **Figure S2D-L**).

### Subcortical responses depend on corticothalamic feedback

To better understand what information is sent back through the extensive descending corticofugal projections to thalamus and midbrain, we optogenetically silenced auditory cortex while recording the responses of an additional 749 units (19 mice) in the IC (*n*=559) and MGB (*n*=190). We tested the effects of cortical silencing on the population communication between the auditory midbrain and thalamus. As expected, auditory cortical silencing significantly decreased the overall firing rate of MGB and IC neurons (although the effect in the IC was small; 0.6% decrease in IC, 29.7% decrease in MGB) (**Figure 4A**, lower panel), demonstrating that descending A1 connectivity modulates the firing rate of the subcortical stations from which it receives its ascending information. We also found that cortical silencing significantly increased the trial-to-trial reliability of units in both IC and MGB (although the effect in IC was again small) (**Figure 4B,E**), suggesting that cortex provides input to subcortical stations which varies from trial to trial and is independent of the spectrotemporal structure of the stimulus.

**Figure 4:**
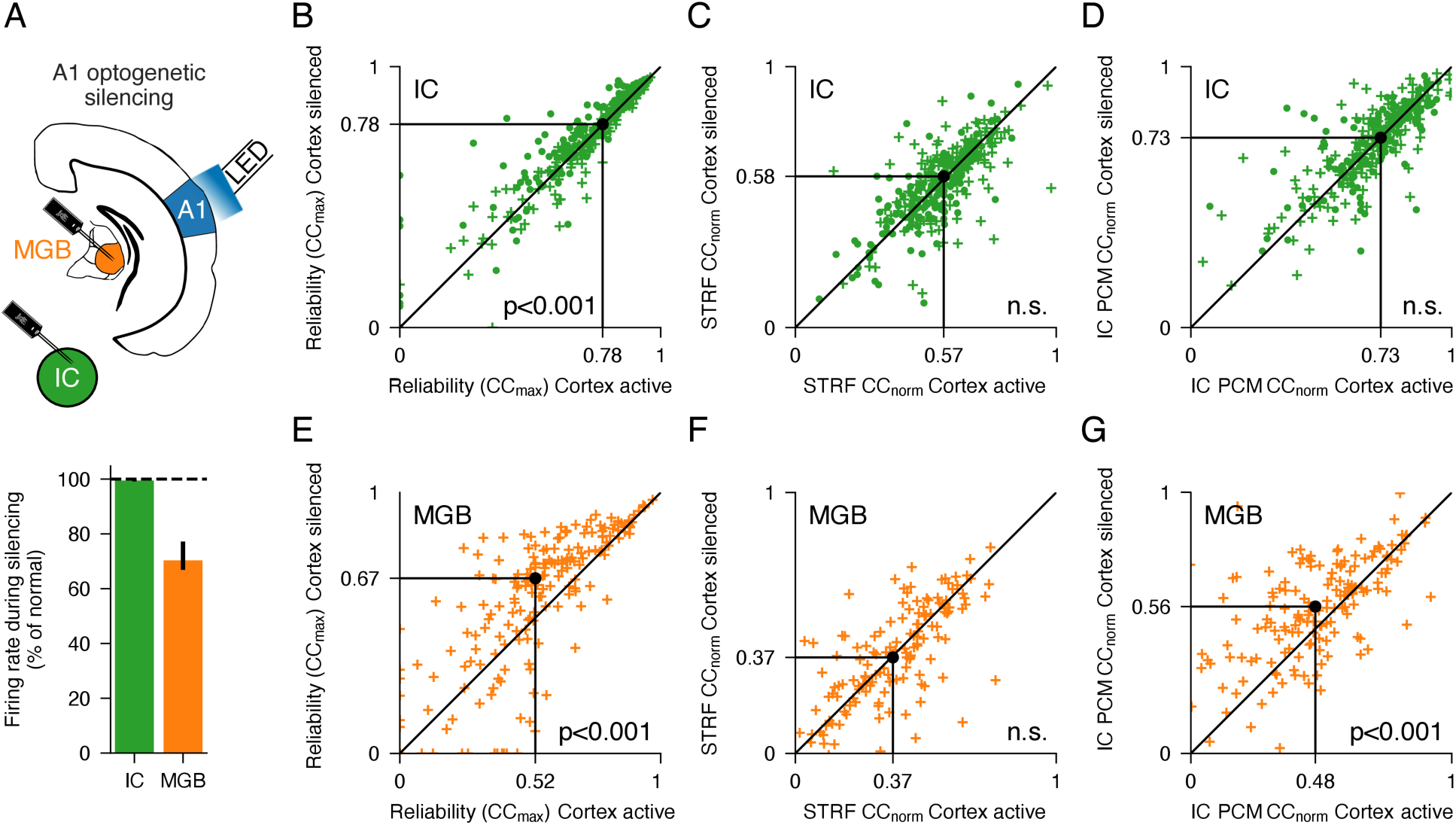
Corticofugal feedback selectively modulates nonlinear responses in MGB. (A) *Top* Schematic of optogenetic silencing experiments. *Bottom* Cortical silencing decreases neuronal firing rate in IC and particularly MGB. Error bars are 95% nonparametric confidence intervals. (B) Trial-to-trial reliability (CC_max_) of IC units with and without auditory cortical silencing. Black dot indicates medians. (C) STRF prediction accuracy (CC_norm_) of IC units with and without auditory cortical silencing. (D) IC PCM prediction accuracy (CC_norm_) of IC units with and without auditory cortical silencing. (E-G) same as in B-D but for MGB.

### Cortical modulation of nonlinear sound encoding in thalamus

We found that STRF model prediction performance was not affected by auditory cortical silencing, after accounting for the change in trial-to-trial reliability (**Figure 4C,F**, *p* > 0.05, *n*_IC_*=* 190, *n*_MGB_ *=* 559). Crucially, however, the IC to MGB population communication model performed significantly better after the cortex was silenced (**Figure 4D,G**, *p* < 0.001, *n*_IC_ *=* 190, *n*_MGB_ *=* 559). This suggests that corticothalamic projections modulate nonlinear stimulus-locked activity in MGB, while leaving the linear representation of spectrotemporal information intact.

Together, these findings reveal that the cortex can influence the activity of thalamic neurons in two distinct ways.

### Cortical control of functional connectivity within subcortical areas

To further explore how descending corticofugal connections control the structure of subcortical activity, we constructed an encoding model of the moment-to-moment spiking activity from the stimulus inputs with (or without) spike coupling between simultaneously-recorded units^31^ (**Figure S7A**). We fitted this model to neural populations recorded in the midbrain and thalamus, and investigated how functional spike coupling between subcortical units was affected by optogenetic silencing of cortex.

We found that including coupling between units in this model increased the predictability of moment-to-moment spiking activity, implying that these responses depend not only on the stimulus, but also on the functional connectivity between neurons in thalamus and midbrain (**Figure 5A-D**, with cortex active or silenced; for each condition, p << 0.001, *n*_IC_*=* 190, *n*_MGB_ *=* 559). Silencing auditory cortex also significantly decreased the contribution of local functional connectivity in explaining both thalamic and midbrain activity, with much stronger effects for the thalamus (**Figure 5E**, *p* < 0.001, *n*_IC_*=* 190, *n*_MGB_ *=* 559, **Figure S7**). This suggests that the descending projections from cortex to thalamus and midbrain allows the orchestration of functional connectivity between neurons within these circuits.

**Figure 5:**
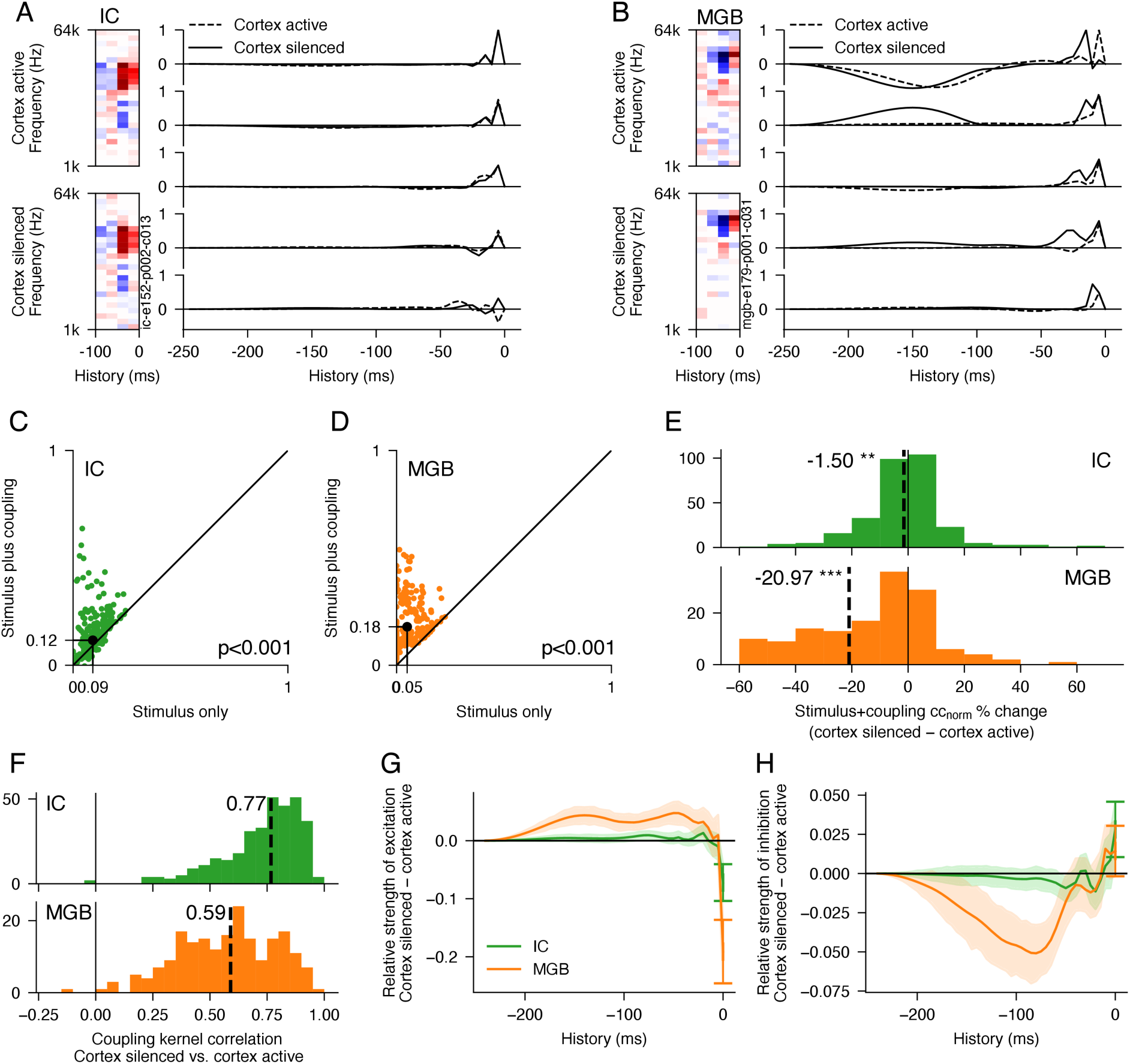
Cortex controls functional spike coupling between simultaneously-recorded neurons in MGB and IC. (A) Example IC unit STRF (*left*) and coupling filters of five simultaneously-recorded IC units (*right*) with and without optogenetic cortical silencing. (B) Same as in A, but for MGB. In each case, these filters were estimated from a single-trial spike coupling GLM model. (C) Comparison of stimulus-only model vs model that also incorporated coupling between simultaneously-recorded IC units in predicting single-trial spike rates. Black dot indicates medians. (D) Same as in c, but for MGB units. (E) Coupling of simultaneously-recorded units adds more explanatory power for IC and especially MGB units when cortex is intact than when cortex is optogenetically silenced. (F) Coupling filters from simultaneously-recorded units change significantly in IC and especially in MGB when cortex is optogenetically silenced. (G) Change in strength of excitatory components of coupling filters in IC and MGB over time after silencing auditory cortex. (H) Change in strength of inhibitory components of coupling filters in IC and MGB over time after silencing auditory cortex. Error bars are 95% confidence intervals.

We also found that the structure of the functional coupling between neurons in IC and MGB was altered when cortex was silenced, with larger changes in MGB than in IC (**Figure 5F**, *p* < 0.001, *n*_IC_*=* 190, *n*_MGB_ *=* 559). In particular, cortical inactivation decreased the fast (<10 ms) excitatory coupling between neurons in thalamus (and to a smaller but significant degree in the midbrain), while increasing the slower (>10 ms) excitatory coupling (**Figure 5G)**. Conversely, fast inhibitory coupling in thalamus was slightly decreased, whereas slower inhibitory coupling was strengthened when cortex was active (**Figure 5H)**. This dynamic cortical control of functional coupling between neurons in the thalamus is consistent with a disruptive effect of silencing cortex on thalamocorticothalamic (excitatory) and thalamocortico-reticulothalamic (inhibitory) open loops between thalamic cells.

## Discussion

We have shown that the representation of complex sound in the auditory cortex arises because of nonlinear transformations between the cochlea and the auditory midbrain, with the midbrain representation then communicated to thalamic neurons, where further nonlinear processing takes place. These subcortical transformations, together with local cortical processing, produce a highly nonlinear representation of sound in the auditory cortex. Importantly, the transformations that occur along the ascending auditory pathway are irreversible, meaning that subcortical responses cannot be reconstructed from the cortical representation that they give rise to.

Although it is well known that spectrotemporal models fail to capture much of the stimulus-locked response variance of auditory cortical neurons^32,33^, it has proven difficult to improve these models in ways that capture this unexplained variance. Network receptive field models modestly outperform simple STRF models for natural sound stimuli^18^ and perform similarly to STRF models for the complex spectrally-random stimuli used here. This has led to the conclusion that much of the unexplained variance in cortical responses may be the result of extra-sensory factors. There has therefore been considerable focus in recent years on how behavioural task demands and brain state differences shape the sensitivity and spectrotemporal tuning properties of auditory cortical neurons^8,25,34–36^.

Here, we show that much of the unexplained variance reflects the failure of existing spectrotemporal models to adequately account for the non-linear spectrotemporal tuning properties of cortical neurons. Population communication models are able to explain 70% more cortical response variance than spectrotemporal models, an improvement that is substantially larger than that achieved by attempts to incorporate nonlinearities into spectrotemporal models^9,12,13,17,18^. This also indicates that a substantial portion of the previously unexplained variance is genuinely stimulus-locked and predictable, and must therefore reflect the structure of auditory stimuli. However, the additional variance cannot be accurately expressed by linear, linear-nonlinear, or network-based spectrotemporal models, suggesting that it reflects fundamentally higher-order representations of the spectrotemporal input. Moreover, by demonstrating that we can predict the responses of A1 neurons to complex acoustic stimuli with considerably more accuracy than has hitherto been possible, our findings indicate that the auditory cortex does play a key role in representing sounds, rather than just the sensory and behavioral context in which those sounds occur.

By also building population communication models of descending connectivity, we have shown that A1 representations are higher order and cannot be converted back into subcortical representations by linear or neural network models. This indicates that the transformation of information from subcortical areas to cortex is lossy and irreversible. This raises the possibility that rather than merely constructing a more complex representation of sound, the cortex also actively discards spectrotemporal information that is necessary for representation in subcortical areas, but which may not be needed for higher-level processing.

Such transformations into higher-order representations are akin to what is found between thalamus and cortex in the visual system^20,37,38^, but have been harder to identify in the auditory system. We find that these transformations occur between the midbrain and thalamus, as well as between thalamus and cortex, and it seems likely that they also take place in the auditory brainstem. An interesting question is whether comparable sequential transformations of information also take place in the other sensory systems. Applying the population communication framework introduced here to large-scale recordings from multiple levels of these systems could illuminate whether this is the case.

It is well known that the auditory cortex is the source of massive descending connections to auditory and other subcortical nuclei^23,24,39^. We found that silencing auditory cortex alters the moment-to-moment spike coupling between cells in the thalamus and midbrain, suggesting that descending corticofugal input may control the timing and balance of excitatory and inhibitory functional connectivity in midbrain and thalamic circuits. This would allow for flexible orchestration of subcortical functional connectivity, potentially controlling which neuronal populations are synchronised and which are not, and therefore which stimulus representations are amplified and transmitted more effectively to cortex^40^. Our optogenetic experiments also indicate that corticothalamic modulation selectively affects the higher-order (nonlinear) representations of sound in the thalamus, leaving linear spectrotemporal tuning intact. Such interactions between cortex and thalamus may happen through specific communication subspaces, as recently found between cortical areas in the visual system^41,42^. This exclusive modulation of higher-order response properties suggests that feedback from the auditory cortex principally influences thalamic responses that are most closely related to its own higher-order sound representation.

Our results highlight two approaches that should lead to a greater understanding of information coding in auditory and other sensory systems. The first is to use the modeled responses of subcortical neurons to build better neural network models of neurons in the cortex. The second is to investigate the contribution that the tuning properties of subcortical neurons make to the high performance of population communication models of information transmission between different processing levels. These approaches are likely to yield fundamental insights into the role of subcortical structures in shaping cortical responses, both in the auditory system and other sensory systems.

## Supplementary Information

**Figure S1:**
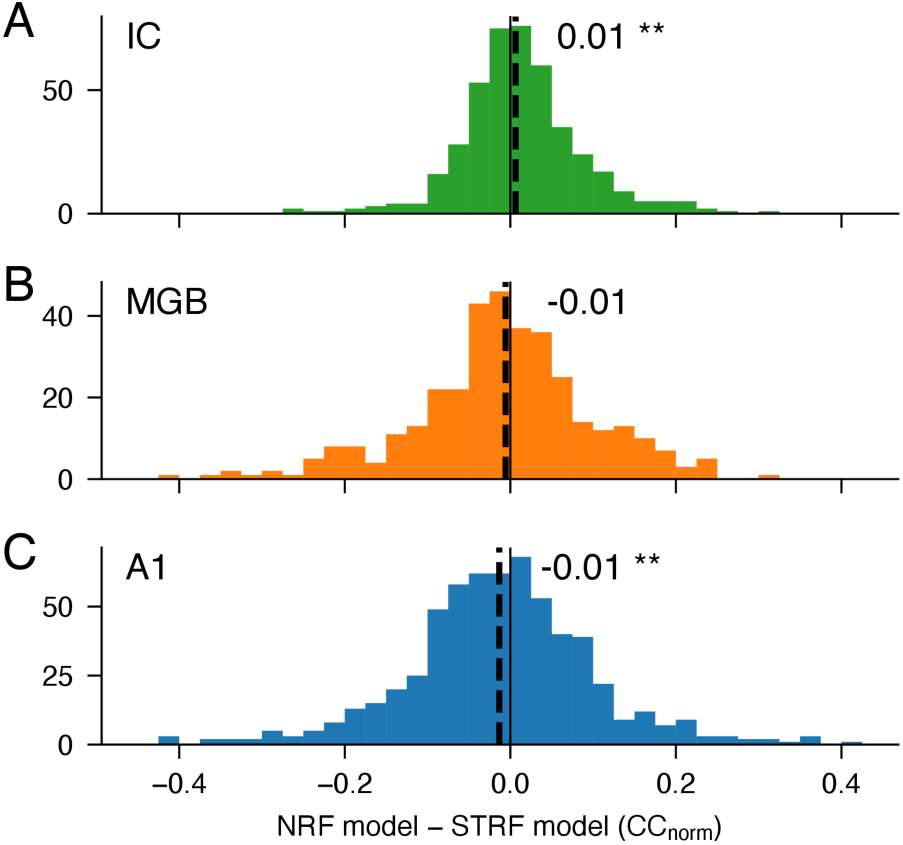
Neural network models perform similarly to linear-nonlinear STRF models on this dataset. (A-C) Relative performance (CC_norm_) of Network Receptive Field (NRF) models and linear-nonlinear STRF models in predicting responses of neurons in IC, MGB, A1 respectively. In this study, we used linear-nonlinear STRF models as our baseline for evaluating the performance of population communication models. It is conceivable, therefore, that the apparently high predictive power of population communication models merely reflects poor performance of our STRF models. To ensure this is not the case, we also fitted NRF models, which describe the responses of each neuron using a neural network whose input is the time-varying cochleagram (log-spectrogram). We find that, for this dataset (where the stimuli are spectrally random), the NRF model outperforms the STRF model for some neurons, and underperforms for others, but there are only minor differences in predictive power across the neural population in all three areas.

**Figure S2:**
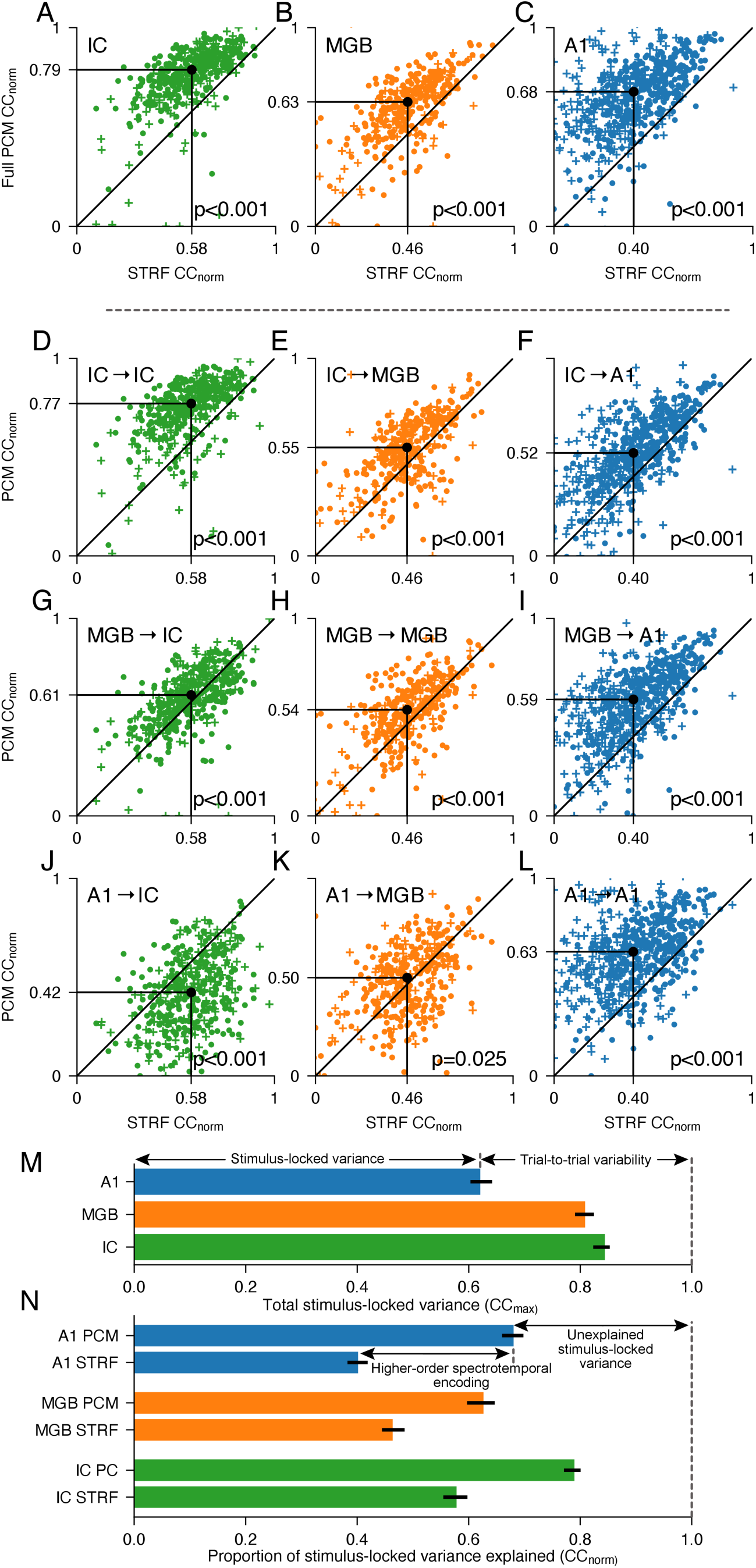
Additional variance explained by the population communication models, compared to STRF models. (A-C) Comparison of prediction performance of full population communication model (taking inputs from non-simultaneously-recorded units in IC, MGB and A1; y-axis), relative to STRF model (x-axis), for predicting responses in IC, MGB and A1 respectively. (D-L) Comparison of prediction performance of single-area population communication models (y-axis), relative to STRF model (x-axis). The source population and target areas are shown in the top-left of each plot (source population → target). (M) Total stimulus-locked variance (cc_max_) in IC, MGB and A1, respectively. The difference from 1 is a measure of how much variance in each area is not stimulus-locked, i.e., trial-to-trial variability in neural responses. (N) Median proportion, cc_norm_, of the stimulus-locked variance (cc_max_) that can be explained by STRF and population communication models for each brain region. The performance of the STRF model is a measure of how much variance can be explained by simple spectrotemporal models. The additional performance of the population communication model indicates the proportion of neuronal responses that can be explained by higher-order spectrotemporal encoding, which is stimulus-locked but cannot be modeled with simple spectrotemporal models. The remaining variance is stimulus-locked, but remains unexplained.

**Figure S3:**
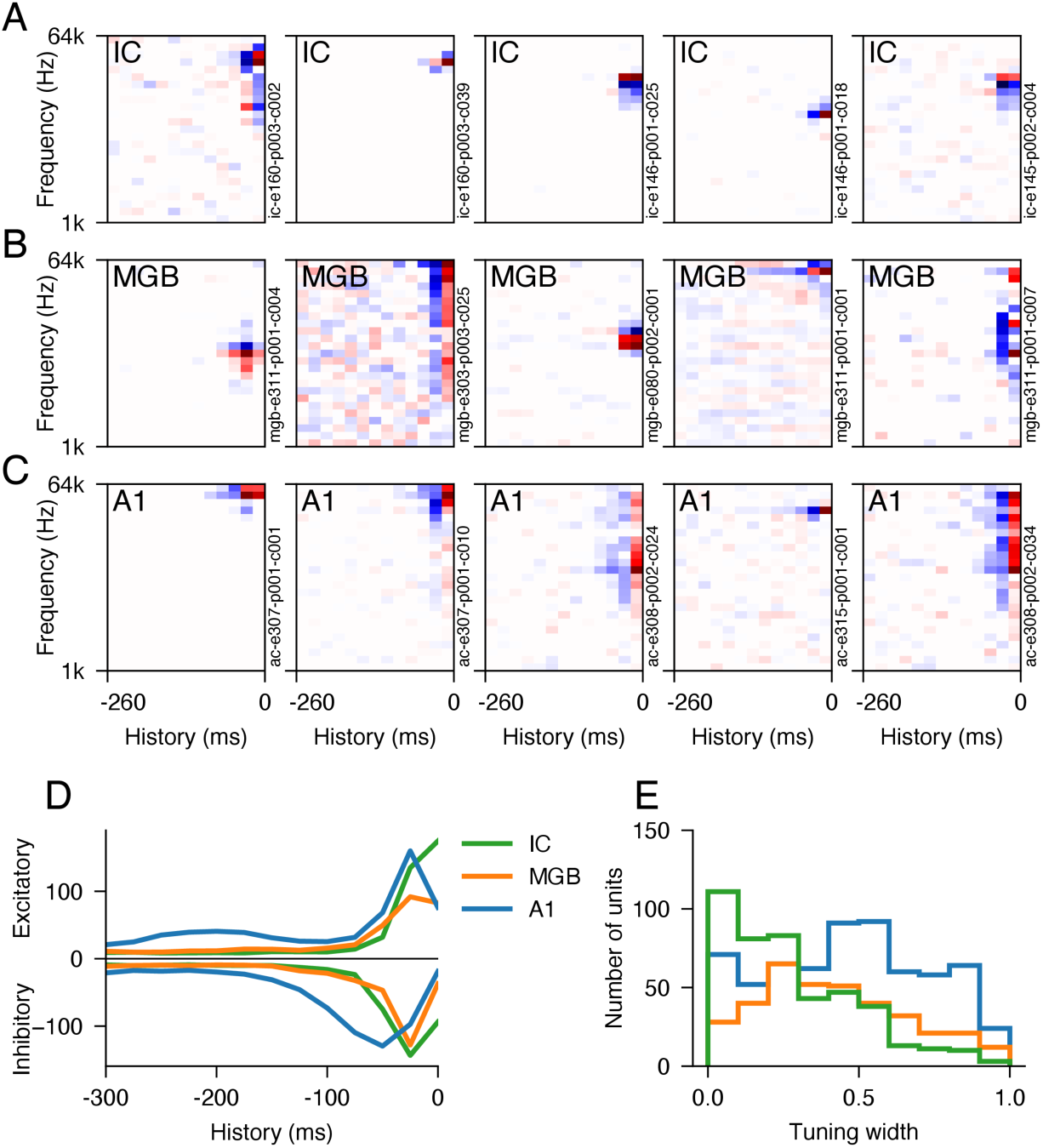
STRF model characteristics over the auditory hierarchy. (A-C) Examples of STRFs in IC, MGB and A1, respectively. (D) Time course of excitatory (top) and inhibitory (bottom) coefficients, summed across all STRFs for each brain area, showing the increase in response latency at higher levels of the auditory hierarchy. (E) Tuning width of STRFs (see Methods), showing an increase across the hierarchy.

**Figure S4:**
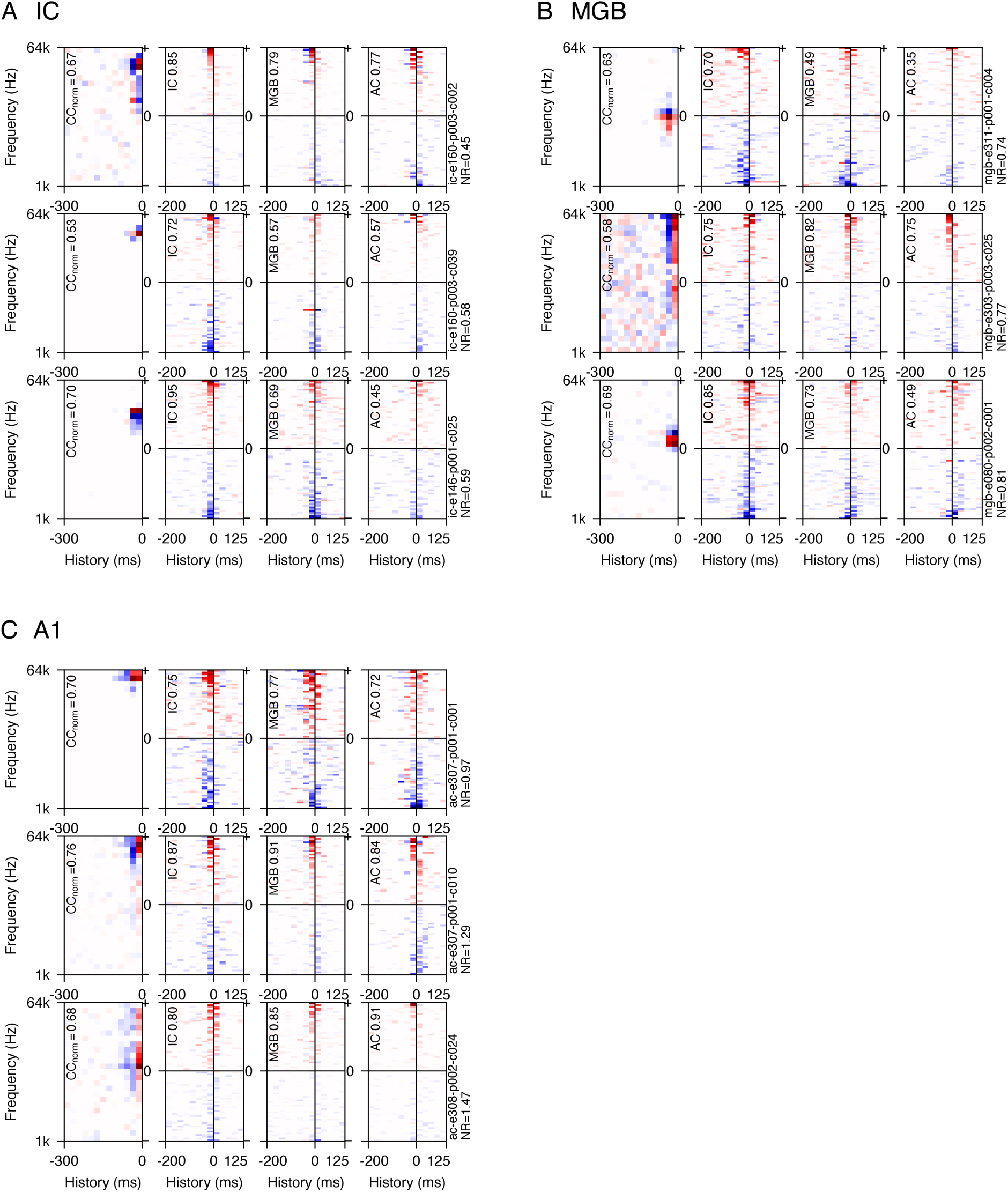
Comparison of STRF and population communication model kernels over the auditory hierarchy. (A) Examples of STRFs and population communication model kernels for three example IC neurons. The left column shows the STRF model kernel, and the subsequent three columns show population communication model kernels using IC, MGB and A1 units as regressors (source population), respectively. CC_norm_ values for each model are shown in the top left. For population communication model kernels, only the regressor units with the 20 highest (above the axis) and 20 lowest (below the axis) summed coefficient values are shown, in descending order of summed coefficient value. (B-C) Similar examples for MGB and A1 units, respectively.

**Figure S5:**
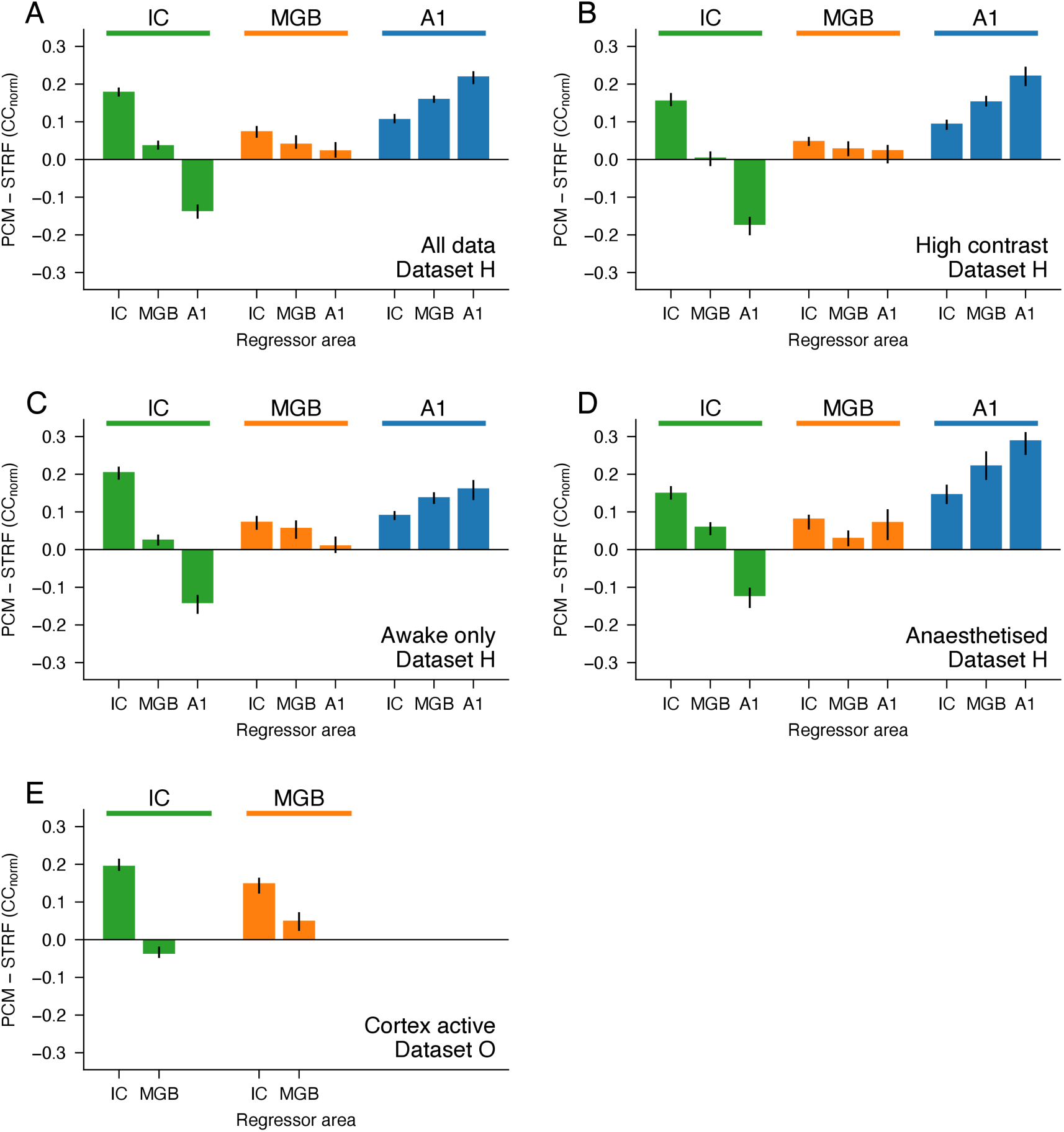
Control analyses on subsets of data. Difference (median) between prediction accuracy of population communication models and STRF models for units recorded in the IC, MGB and A1 (target population) with regressor inputs (source population) from different processing levels. The values in these panels are comparable to those in Figure 3D, but are measured for different subsets of the data. (A) Main results for dataset H, as in Figure 3D. (B) Equivalent analysis for dataset H, but using only neural responses to high contrast sounds. (C) Equivalent analysis including only awake data from dataset H. (D) Equivalent analysis including only anesthetised data from dataset H. (E) Equivalent analysis using dataset O (optogenetic data), including only the condition where cortex was active.

**Figure S6:**
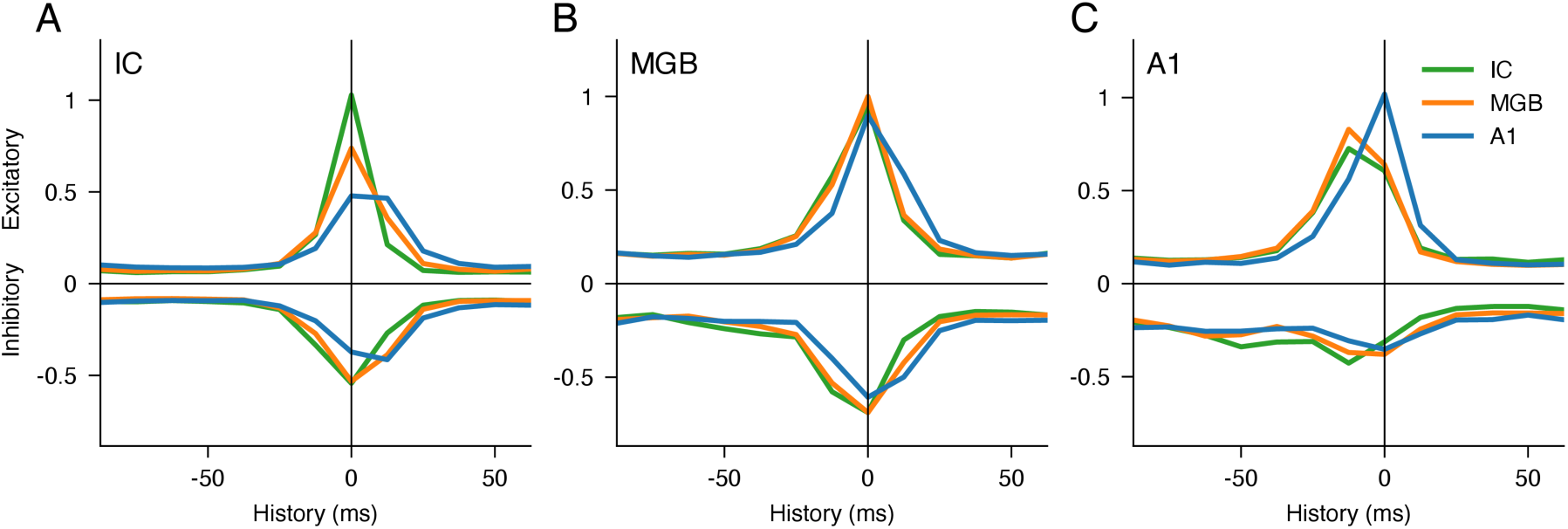
Time course of population communication models over the auditory processing hierarchy. (A) Time course of excitatory (above x-axis) and inhibitory (below x-axis) coefficient values in population communication models of IC neurons, summed over regressor units (source population) in single-area population communication models. The mode offset for IC-to-IC coefficient values is 0, and increases for regressor units located in higher auditory areas (MGB and A1). This is expected, given the increasing latency of typical neural responses across the auditory pathway. (B-C) Similar plots for MGB and A1 target units, respectively.

**Figure S7:**
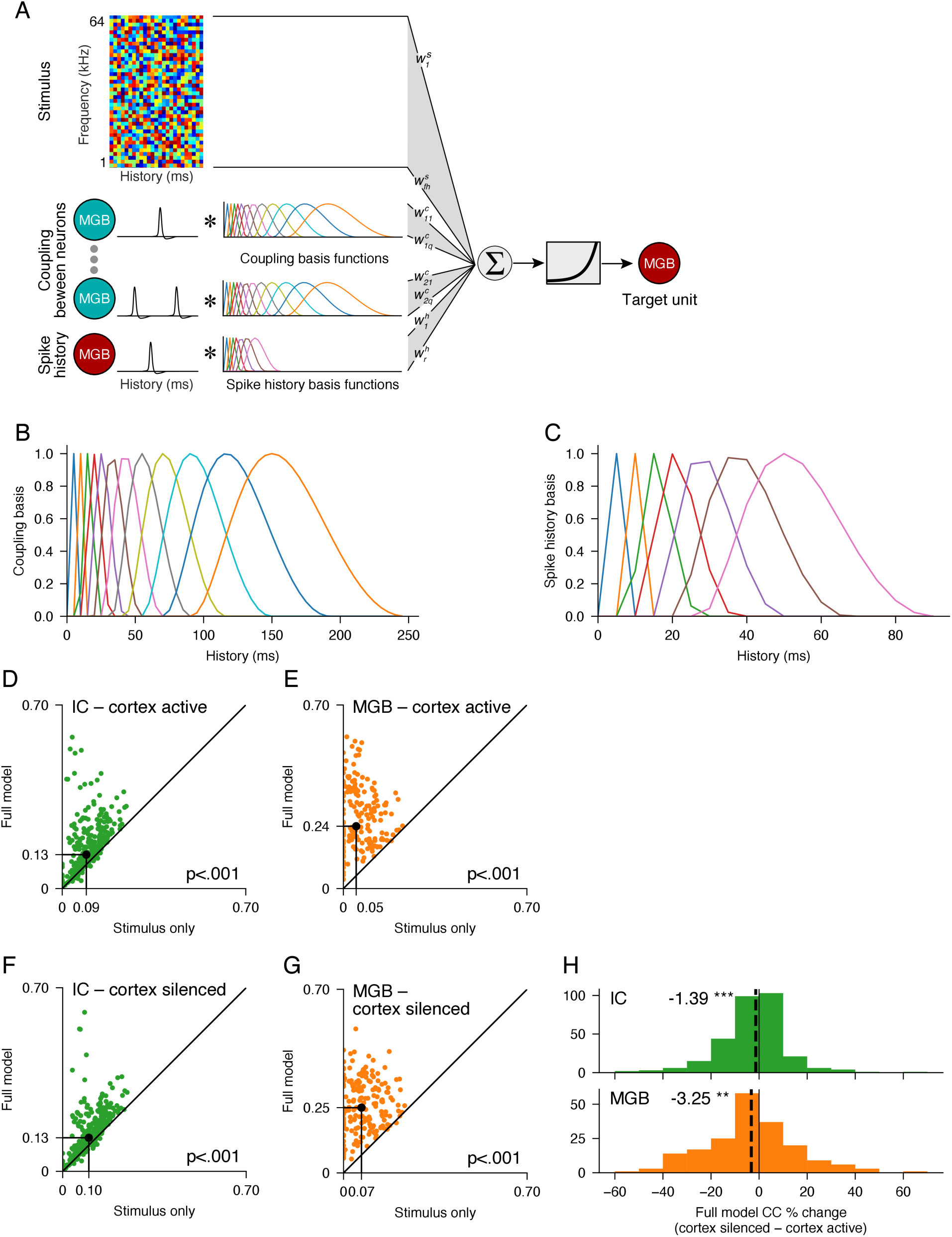
GLM analysis of neuron-to-neuron communication. (A) Schematic showing the full GLM model, which consists of an STRF component (similar to STRF models), with additional regressors describing coupling between simultaneously-recorded neurons and the spike history of the target neuron. The spike coupling regressors are the result of convolving the responses of simultaneously-recorded neurons with basis functions (B) spanning latencies up to 250 ms at 5 ms resolution. The spike history regressors are the result of convolving the target neuron’s responses with basis functions (C) spanning latencies up to 90ms. To fit the neural responses, weights are estimated for all regressors which minimise the MSE between the model output and the target neuron’s responses, using a Poisson GLM^31^. (D-G) The full model, including coupling and spike history, predicts neuronal responses better than the same model including the stimulus only, in both IC (D, F) and MGB (E, G), and both when the cortex was active (D, E) and when it was silenced optogenetically (F, G). (H) Performance of the full model is slightly higher when the cortex was active than when the cortex was silenced, for both IC and MGB.

## Acknowledgements

This work was funded by a Wellcome Doctoral Training studentship (WT105241/Z/14/Z) and Sir Henry Wellcome postdoctoral fellowship (224121/Z/21/Z) to M.L. and a Wellcome Principal Research Fellowship (WT108369/Z/2015/Z) to A.J.K. We are grateful to Mollie Ward and Thomas Blennerhassett for their contributions to the analyses.

## Author contributions

M.L. and A.J.K. conceived and designed the experiments, M.L. performed the experiments, B.D.B.W. and M.L. analysed the data, and all authors wrote, revised and edited the manuscript.

## Code availability

All code used to generate analyses and figures presented in this manuscript will be available at https://github.com/ben-willmore/population-communication-models

## Data availability

Data is available upon request to

## Methods

### Mice

All animal experiments conformed to ethical standards approved by the Committee on Animal Care and Ethical Review at the University of Oxford and were licensed by the UK Home Office (Animal Scientific Procedures Act, 1986, amended in 2012). A total of 47 mice were used in this study. Four strains of male and female mice were used: *C57BL6/J* (Envigo, UK), *GAD2-IRES-cre* (Jackson Laboratories, USA), *VGAT-ChR2-YFP* (Jackson Laboratories, USA), and *C57BL6/NTac.Cdh23*^43^. *C57BL6/J*, *GAD2-IRES-cre*, and *VGAT-ChR2-YFP* were 7–12-weeks old at the time of data collection, and *C57BL6/Ntac.Cdh23* were 10–20-weeks old at the time of data collection. All experiments were carried out in a sound-attenuated chamber.

### Stimuli

Stimuli were presented with a Tucker-Davis Technologies (TDT) RX6 Multifunction processor at ∼200 kHz. Sounds were amplified by a TDT SA1 stereo amplifier and delivered via a modified Avisoft ultrasonic electrostatic loudspeaker (Vifa) positioned ∼1 mm from the ear canal. The sound presentation system was calibrated to a flat (±1 dB) frequency-level response between 500 and 64,000 Hz.

Stimuli consisted of spectrotemporally complex dynamic random chords (DRCs) with individual chords having a duration of 25 ms (including 5 ms on and off ramps) and comprising 25 superposed frequencies, logarithmically spaced between 1000 and 64,000 Hz (1/4 octave intervals). The tones of the DRC were played at sound levels that were randomly drawn from one of two uniform distributions: 30–50 dB sound pressure level (SPL) (low contrast) or 20–60 dB SPL (high contrast). The mean of the distribution was therefore constant, at 40 dB SPL. The logarithmic statistics of the decibel scale have been found to better match the statistics of natural sounds^44,45^. The overall sound level of the DRCs was calibrated to be 79–83 dB SPL. DRCs for any given trial were played for either 40 s (dataset H in **Figure S5**) or 5 s (dataset O in **Figure S5**; 5-s trial duration in optogenetic experiments), with inter-trial intervals of 2–10 s.

### In vivo extracellular recordings

We carried out extracellular recordings using 32- or 64-channel silicon probes (NeuroNexus Technologies Inc.), in a 4 × 8, 8 × 8, or 2 × 32 electrode configuration. Electrophysiological data were acquired on a Tucker-Davis technologies (TDT) RZ2 BioAmp processor and collected and saved using custom-written Matlab code (https://github.com/ben-willmore/benware).

#### Awake recordings

For awake recordings, we chronically implanted a recording chamber under isoflurane (1.5–2% in oxygen) general anaesthesia together with administration of meloxicam (5 mg/kg) and dexamethasone (Dexadreson, 2-3mg/kg). The recording chamber consisted of a metal cylinder positioned over a craniotomy, with a lightly attached circular window in order to close the recording chamber. We placed the recording chamber around craniotomies over IC (centered ∼5 mm posterior from bregma and ∼1 mm lateral from midline), A1 (centered ∼2.5 mm posterior from bregma and ∼4.5 mm lateral from midline), or the cortex above MGB (centered ∼3 mm posterior from bregma and ∼2.1 mm lateral from midline), together with a head bar and a reference (silver wire) in the contralateral hemisphere. We then fixed the implant to the skull using a dental adhesive resin cement (Super Bond C&B). Following full recovery, on a subsequent day the mouse was head-fixed, the recording chamber was opened, and the exposed dura mater was kept moist with saline. A sterile recording probe was then acutely inserted into the into the recording site of interest. All recordings were performed in the right hemisphere.

In the mouse, the dorsal surface of the IC is not covered by the cortex and is very distinct. The craniotomies over the IC were always large enough to see the entire exposed IC surface, so we could visually target the probes. We inserted the probe in the center of the IC, and therefore above the central nucleus of the IC (CNIC), where the overlying dorsal cortex is relatively thin^46^. We confirmed this by checking for the clear dorso-ventral tonotopic gradient in the STRFs that is indicative of this nucleus. We also estimated frequency response areas using tones, which confirmed the presence of dorso-ventral tonotopic gradients with narrow tuning (data not shown). When we were positioning the electrode array, we observed tightly-locked multiunit responses to noise stimuli, characteristic of CNIC neurons, and post-mortem inspection of the midbrain confirmed that the probe had indeed been located in the CNIC.

Prior to insertion into auditory thalamus, the probe was coated in DiI (Sigma-Aldrich) for subsequent histological verification of the recording site. Recording sites were confirmed as being located in auditory thalamus if multiunit activity responded to broadband noise and was frequency tuned when the tip of the probe was ∼2.5–3.5 mm below the brain surface. Auditory thalamic recordings were subsequently attributed to the lemniscal ventral subdivision of the MGB (MGBv) by histological investigation of recording sites. Based on an immunohistochemical study by Lu et al. (2009)^47^ on the shape and size of subdivisions of the mouse auditory thalamus, we allocated recording sites to the MGBv if they responded reliably to DRC stimulation on electrode channels < 500 μm from the lateral border of the MGB.

Finally, A1 was identified by robust neuronal responses to broadband noise bursts, and a caudo-rostral tonotopic axis.

#### Anaesthetized recordings

For experiments carried out under anaesthesia, mice were anaesthetized with an intraperitoneal injection of ketamine (100 mg kg^−1^) and medetomidine (0.14 mg kg^−1^). We also administered intraperitoneal injections of atropine (Atrocare, 1 mg kg^−1^) to prevent bradycardia and reduce bronchial secretions, and dexamethasone (4 mg kg^−1^) to prevent brain edema. Prior to initial surgery, bupivacain was administered as an analgesic under the scalp. The depth of anaesthesia was monitored via the pedal reflex and small additional doses of the ketamine/medetomidine mix were given subcutaneously approximately every 15 min once the recordings started (∼1–1.5 h post induction of anaesthesia). The dosage of individual top-ups depended on the depth of anaesthesia at the time, but corresponded to ∼50 mg kg^−1^ h^−1^ of ketamine and ∼0.07 mg kg^−1^ h^−1^ of medetomidine. All recordings were performed in the right hemisphere. A silver reference wire was positioned in the visual cortex of the contralateral hemisphere, and a grounding wire was attached under the skin on the neck. The head was fixed in position with a metal bar acutely attached with bone cement to the skull over the left hemisphere. We then made 2-mm diameter circular craniotomies above the IC (centred ∼5 mm posterior from bregma and ∼1 mm lateral from midline), over the visual cortex for auditory thalamic recordings (centred ∼3 mm posterior from bregma and ∼2.1 mm lateral from midline), and/or over the auditory cortex (centred ∼2.5 mm posterior from bregma and ∼4.5 mm lateral from midline). Following exposure of the brain, the exposed dura mater was kept moist with saline. The silicon probe was then inserted carefully into the recording site of interest.

### Optogenetic silencing of auditory cortex

The data for the optogenetic silencing experiments were also used in Lohse et al. (2020)^14^, where results confirming electrophysiological effective silencing of activity are reported.

To transiently silence the activity of auditory cortical excitatory neurons, we employed either a transgenic or a viral approach to express ChR2 in auditory cortical inhibitory neurons. *VGAT-ChR2-YFP* mice express ChR2-YFP in GABAergic neurons throughout the adult brain. Optogenetic activation of cortical inhibitory neurons is the most effective available method for inhibiting cortical activity at sub-second time resolution over the time window required for this study and has been used extensively to transiently silence excitatory activity (including corticofugal outputs) in cortical areas in mice. Viral injection surgeries were performed under isoflurane (∼1.5%) anaesthesia, with the animal positioned in a stereotaxic frame (Kopf instruments, USA). For viral transfection, we injected *AAV5-DIO-ChR2-eYFP* (UNC gene therapy vector core) into the auditory cortex of *GAD2-IRES-cre* mice. We injected ∼400 nl of virus, spread over three locations (spaced caudal-rostrally ∼400 μm apart) at three depths (700, 500, and 300 μm from cortical surface), to ensure widespread expression in auditory cortex. Mice were used for electrophysiological recordings > 4 weeks post injection of virus. This ensured strong expression of ChR2-eYFP in the auditory cortex.

For optogenetic silencing, we exposed the auditory cortex to blue (470 nm) LED light. This was achieved by placement of a 200 μm (*VGAT-ChR2-YFP* experiments) or 1 mm optical fibre (*GAD2-cre* + viral ChR2 experiments) immediately above the dura mater over the auditory cortex to allow for blue light exposure to ChR2-expressing cells. For silencing of auditory cortical activity during recordings in MGBv or CNIC, we stimulated with blue light at 40 Hz frequency using sinusoidal waves or 15 ms pulses (10 ms gaps). When recording from auditory cortex, we stimulated with blue light at 40 Hz using either sinusoidal waves or 15 ms pulses (10 ms gaps) or constant light stimulation. Light power was ∼5–7 mW mm^−2^ at the tip of the fiber. We found that light stimulation (40 Hz (sinusoid or pulsed) or constant light) effectively silenced activity in auditory cortical neurons by driving inhibitory neurons for the duration of the DRC stimulation (5 s) (see Lohse et al., 2020^14^).

### Histology

For post-mortem verification of the electrophysiological recording sites and viral expression patterns, mice were overdosed with pentobarbital (100 mg/kg body weight, i.p.; pentobarbitone sodium; Merial Animal Health Ltd, Harlow, UK) and perfused transcardially, first with 0.1 M phosphate-buffered saline (PBS, pH 7.4) and then with fresh 4% paraformaldehyde (PFA, weight/volume) in PBS.

### Spike sorting

We clustered potential neuronal spikes using KiloSort (https://github.com/cortex-lab/KiloSort). Following this automatic clustering step, we manually inspected the clusters in Phy (https://github.com/kwikteam/phy), and removed noise (movement artefacts, optogenetic light artefacts etc.). We assessed clusters according to suggested guidelines published by Stephen Lenzi and Nick Steinmetz (https://phy-contrib.readthedocs.io/en/latest/template-gui/#user-guide).

### Modeling

After spike sorting, data collation was conducted in MATLAB, after which data was analyzed using Python code based on NumPy, scikit-learn and glmnet-python (https://github.com/bbalasub1/glmnet_python/tree/master/glmnet_python).

#### Neural responses

For each unit, we counted spikes in 25 ms time bins for the STRF and population communication models (corresponding to the chord length of the DRC stimuli), giving matrix *y_dt_*. For the single trial poisson Generalized Linear Model, spikes were counted in 5ms bins. Where appropriate, we averaged these counts over all trials to compute the peristimulus time histogram (PSTH), resulting in vector *y_t_*.

#### Unit selection criteria

We aimed to include as many units as possible in our analyses, excluding only units where responses were very unreliable (likely corresponding to poorly isolated clusters). To achieve this, we measured reliability using the noise ratio^10,48^ (NR) of the responses, *y_dt_*, of each unit. Each unit was included in our analyses if it had a noise ratio of < 200 across the entire dataset. In order to make unbiased comparisons between different models, we also excluded from all analyses in **Figures 1-3** any unit where any model failed to produce a fit. For **Figures 1-3**, this meant that 120 out of 1561 recorded units were excluded (giving a total of 1441 included units). For optogenetic recording data related to **Figures 4 and 5**, 44 out of 793 units were excluded (giving a total of 749 included units).

#### Testing on a held-out data set

To fit and test the STRF and population communication models, we divided the stimulus into *n* equally sized contiguous segments (*n=*16 for **Figures 1-3**, *n=*15 for **Figures 4 and 5**). For every fit, models were trained on neuronal responses to the first (*n*-1) segments, and hyperparameter selection was carried out on a subset of the first (*n*-1) segments. Models were tested on the remaining (*n*^th^) segment, so all correlation coefficient values are for prediction on a “held-out” dataset, i.e., a subset that had not been used in any part of the training procedure. This approach means that the reported correlation coefficients are resistant to overfitting – i.e., models with greater numbers of regressors do not have a built-in advantage over models with fewer regressors. Due to the computation time required, we did not fit every model on all *n* folds of the data, though our main results were validated on all folds.

For GLM fitting, where trial-to-trial variability is included, models were trained on 5 concatenated single-trial responses to the entire stimulus. A held-out test set of 2 concatenated single-trial responses to the entire stimulus was used for model evaluation.

#### Model evaluation

The prediction performance of STRF and population communication models was evaluated on the held-out test set using normalized correlation coefficient^28^ (cc_norm_), i.e. the correlation coefficient between the predicted and actual responses after estimated trial-to-trial variability has been excluded. For GLM models, trial-to-trial variability is of interest and cannot be excluded. For these models, prediction performance was evaluated on the held-out test set using the standard Pearson correlation coefficient.

#### Linear-nonlinear STRF models

We characterized the power in each stimulus using the raw sound-level (dB SPL) vs time values from which the DRCs were originally created. These values constitute a log-spectrogram (referred to as a cochleagram), *X_tf_*, where *t* indexes time in 25 ms steps, and *f* indexes the stimulus frequencies. The spectrogram contains complete information about the time-varying frequency content of the stimulus. We ‘tensorized’ the stimulus matrix by adding an extra dimension containing the unrolled stimulus history (over the most recent 13 time steps) at each time point, resulting in a spectrogram tensor, *X_tfh_*, where *h* indexes stimulus history. We then estimated a linear model consisting of a spectrotemporal kernel, *k_fh_*, and offset, *a*_0_, which is applied to the spectrogram tensor to produce a linear estimate, *z_t_*, of the time-varying neuronal responses:

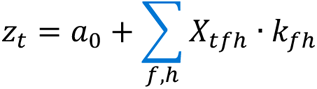

The values of *k_fh_* and *a* were optimized using scikit-learn to minimize the mean square error between *z_t_* and *y_t_*, subject to L2 regularization. Regularization strength was determined by a hyperparameter, *λ*, whose value was selected by cross-validation on a held-out subset of the training data (not overlapping with the test set). Finally, we applied a logistic output nonlinearity, which transforms the linear model output, giving the final modelled neuronal responses 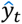:

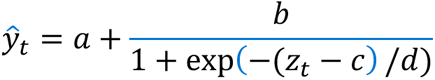

Parameters *a*, *b*, *c* and *d* were optimized by gradient descent to minimize mean-square error between 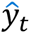 and *y_t_*.

#### Linear-nonlinear population communication models

The linear-nonlinear population communication models were identical in form and optimization to the linear-nonlinear STRF models and differed only in the inputs to the regression procedure. Thus, these models consisted of a linear component which described the mean PSTH, *y_t_*, in terms of the tensorized PSTHs of the input neurons, *X_tuh_*, where *u* indexes neuron number. This was followed by a logistic output nonlinearity, as above. To allow for differences in neural response latency, the response history of input neurons was tensorized over 8 history steps and 5 future steps.

#### Network Receptive Field models

For Network Receptive Field (NRF) models^18^, stimuli and neuronal responses were processed exactly as for the linear-nonlinear models. The network model (built using PyTorch) contained a single hidden layer of 2, 4, 8 or 16 hidden units with logistic activation functions, and an output unit with logistic activation function. The number of hidden units was treated as a hyperparameter which was chosen based on prediction performance (mean square error) on a held-out subset of the training data (not overlapping with the test set).

#### Single Trial Poisson Generalized Linear Model with Spike Coupling

The single trial poisson Generalized Linear Model (GLM) analyses used a Poisson GLM with dependence on stimulus and spike history, as well as coupling between neurons^31,49^. To capture stimulus dependence, the design matrix contained the sound level history of the most recent 5 chords (125ms) at 5ms resolution. To capture coupling between neurons, we convolved the responses of each simultaneously-recorded neuron with a set of 12 basis functions spanning 250ms (**Figure S7B**; delta functions at latencies 5ms and 10ms, followed by logarithmically-spaced raised cosine filters; peak latency of longest filter = 150ms). Finally, to capture spike history, we convolved the spike history of the neuron with a set of 7 basis functions spanning 100ms (**Figure S7C**; delta functions at latencies 5ms and 10ms, followed by logarithmically-spaced raised cosine filters; peak latency of longest filter = 50ms). GLMs were fitted using glmnet-python. The fitted coupling and spike history filters were reconstructed by multiplying each filter by the corresponding fitted coefficient, resulting in a time-varying filter relating the target neuron’s response to the recent history of each regressor units’ spiking activity.

### Statistical inference

Unless specified otherwise, all *p*-values were estimated using non-parametric Wilcoxon signed-rank tests (paired samples) or Mann-Whitney U tests (independent samples). Ninety-five percent non-parametric error bars were estimated using sampling with replacement to obtain a distribution of bootstrapped median values, and from this distribution the 2.5 and 97.5 percentiles were identified to create 95% confidence intervals around the median.

## References

1. Kim, P.J., and Young, E.D. (1994). Comparative analysis of spectro-temporal receptive fields, reverse correlation functions, and frequency tuning curves of auditory-nerve fibers. J Acoust Soc Am 95, 410–422. 10.1121/1.408335.

2. Chechik, G., and Nelken, I. (2012). Auditory abstraction from spectro-temporal features to coding auditory entities. Proceedings of the National Academy of Sciences 109, 18968– 18973. 10.1073/pnas.1111242109.

3. Carandini, M. (2006). What simple and complex cells compute. J Physiol 577, 463–466. 10.1113/jphysiol.2006.118976.

4. Harris, K.D., and Mrsic-Flogel, T.D. (2013). Cortical connectivity and sensory coding. Nature 503, 51–58. 10.1038/nature12654.

5. Kato, H.K., Asinof, S.K., and Isaacson, J.S. (2017). Network-level control of frequency tuning in auditory cortex. Neuron 95, 412–423.e4. 10.1016/j.neuron.2017.06.019.

6. Wood, K.C., Blackwell, J.M., and Geffen, M.N. (2017). Cortical inhibitory interneurons control sensory processing. Curr Opin Neurobiol 46, 200–207. 10.1016/j.conb.2017.08.018.

7. Ohzawa, I., Sclar, G., and Freeman, R.D. (1985). Contrast gain control in the cat’s visual system. J Neurophysiol 54, 651–667. 10.1152/jn.1985.54.3.651.

8. Fritz, J., Shamma, S., Elhilali, M., and Klein, D. (2003). Rapid task-related plasticity of spectrotemporal receptive fields in primary auditory cortex. Nat Neurosci 6, 1216–1223. 10.1038/nn1141.

9. Ahrens, M.B., Linden, J.F., and Sahani, M. (2008). Nonlinearities and contextual influences in auditory cortical responses modeled with multilinear spectrotemporal methods. J Neurosci 28, 1929–1942. 10.1523/JNEUROSCI.3377-07.2008.

10. Rabinowitz, N.C., Willmore, B.D.B., Schnupp, J.W.H., and King, A.J. (2011). Contrast gain control in auditory cortex. Neuron 70, 1178–1191. 10.1016/j.neuron.2011.04.030.

11. King, J.L., Lowe, M.P., Stover, K.R., Wong, A.A., and Crowder, N.A. (2016). Adaptive processes in thalamus and cortex revealed by silencing of primary visual cortex during contrast adaptation. Current Biology 26, 1295–1300. 10.1016/j.cub.2016.03.018.

12. Williamson, R.S., Ahrens, M.B., Linden, J.F., and Sahani, M. (2016). Input-specific gain modulation by local sensory context shapes cortical and thalamic responses to complex sounds. Neuron 91, 467–481. 10.1016/j.neuron.2016.05.041.

13. David, S. V. (2018). Incorporating behavioral and sensory context into spectro-temporal models of auditory encoding. Hear Res 360, 107–123. 10.1016/j.heares.2017.12.021.

14. Lohse, M., Bajo, V.M., King, A.J., and Willmore, B.D.B. (2020). Neural circuits underlying auditory contrast gain control and their perceptual implications. Nat Commun 11, 324. 10.1038/s41467-019-14163-5.

15. Machens, C.K., Wehr, M.S., and Zador, A.M. (2004). Linearity of cortical receptive fields measured with natural sounds. J Neurosci 24, 1089–1100. 10.1523/JNEUROSCI.4445-03.2004.

16. Atencio, C.A., Sharpee, T.O., and Schreiner, C.E. (2008). Cooperative nonlinearities in auditory cortical neurons. Neuron 58, 956–966. 10.1016/j.neuron.2008.04.026.

17. Rabinowitz, N.C., Willmore, B.D.B., Schnupp, J.W.H., and King, A.J. (2012). Spectrotemporal contrast kernels for neurons in primary auditory cortex. J Neurosci 32, 11271–11284. 10.1523/JNEUROSCI.1715-12.2012.

18. Harper, N.S., Schoppe, O., Willmore, B.D.B., Cui, Z., Schnupp, J.W.H., and King, A.J. (2016). Network receptive field modeling reveals extensive integration and multi-feature selectivity in auditory cortical neurons. PLoS Comput Biol 12, e1005113. 10.1371/journal.pcbi.1005113.

19. Escabí, M.A., and Read, H.L. (2003). Representation of spectrotemporal sound information in the ascending auditory pathway. Biol Cybern 89, 350–362. 10.1007/s00422-003-0440-8.

20. Priebe, N.J., and Ferster, D. (2012). Mechanisms of neuronal computation in mammalian visual cortex. Neuron 75, 194–208. 10.1016/j.neuron.2012.06.011.

21. Theunissen, F.E., and Elie, J.E. (2014). Neural processing of natural sounds. Nat Rev Neurosci 15, 355–366. 10.1038/nrn3731.

22. Suga, N. (2012). Tuning shifts of the auditory system by corticocortical and corticofugal projections and conditioning. Neurosci Biobehav Rev 36, 969–988. 10.1016/j.neubiorev.2011.11.006.

23. Bajo, V.M., and King, A.J. (2013). Cortical modulation of auditory processing in the midbrain. Front Neural Circuits 6, 114. 10.3389/fncir.2012.00114.

24. Souffi, S., Nodal, F.R., Bajo, V.M., and Edeline, J.-M. (2021). When and how does the auditory cortex influence subcortical auditory structures? New insights about the roles of descending cortical projections. Front Neurosci 15, 690223. 10.3389/fnins.2021.690223.

25. Guo, W., Clause, A.R., Barth-Maron, A., and Polley, D.B. (2017). A corticothalamic circuit for dynamic switching between feature detection and discrimination. Neuron 95, 180–194.e5. 10.1016/j.neuron.2017.05.019.

26. Homma, N.Y., Happel, M.F.K., Nodal, F.R., Ohl, F.W., King, A.J., and Bajo, V.M. (2017). A role for auditory corticothalamic feedback in the perception of complex sounds. J Neurosci 37, 6149–6161. 10.1523/JNEUROSCI.0397-17.2017.

27. Bajo, V.M., Nodal, F.R., Moore, D.R., and King, A.J. (2010). The descending corticocollicular pathway mediates learning-induced auditory plasticity. Nat Neurosci 13, 253–260. 10.1038/nn.2466.

28. Schoppe, O., Harper, N.S., Willmore, B.D.B., King, A.J., and Schnupp, J.W.H. (2016). Measuring the performance of neural models. Front Comput Neurosci 10, 10. 10.3389/fncom.2016.00010.

29. Nelken, I., Fishbach, A., Las, L., Ulanovsky, N., and Farkas, D. (2003). Primary auditory cortex of cats: feature detection or something else? Biol Cybern 89, 397–406. 10.1007/s00422-003-0445-3.

30. King, A.J., and Nelken, I. (2009). Unraveling the principles of auditory cortical processing: can we learn from the visual system? Nat Neurosci 12, 698–701. 10.1038/nn.2308.

31. Pillow, J.W., Shlens, J., Paninski, L., Sher, A., Litke, A.M., Chichilnisky, E.J., and Simoncelli, E.P. (2008). Spatio-temporal correlations and visual signalling in a complete neuronal population. Nature 454, 995–999. 10.1038/nature07140.

32. Theunissen, F.E., Sen, K., and Doupe, A.J. (2000). Spectral-temporal receptive fields of nonlinear auditory neurons obtained using natural sounds. J Neurosci 20, 2315–2331. 10.1523/JNEUROSCI.20-06-02315.2000.

33. Gourévitch, B., Noreña, A., Shaw, G., and Eggermont, J.J. (2009). Spectrotemporal receptive fields in anesthetized cat primary auditory cortex are context dependent. Cerebral Cortex 19, 1448–1461. 10.1093/cercor/bhn184.

34. Schneider, D.M., Sundararajan, J., and Mooney, R. (2018). A cortical filter that learns to suppress the acoustic consequences of movement. Nature 561, 391–395. 10.1038/s41586-018-0520-5.

35. Schwartz, Z.P., Buran, B.N., and David, S. V. (2020). Pupil-associated states modulate excitability but not stimulus selectivity in primary auditory cortex. J Neurophysiol 123, 191–208. 10.1152/jn.00595.2019.

36. De Franceschi, G., and Barkat, T.R. (2021). Task-induced modulations of neuronal activity along the auditory pathway. Cell Rep 37, 110115. 10.1016/j.celrep.2021.110115.

37. Hubel, D.H., and Wiesel, T.N. (1962). Receptive fields, binocular interaction and functional architecture in the cat’s visual cortex. J Physiol 160, 106–154. 10.1113/jphysiol.1962.sp006837.

38. Ferster, D., Chung, S., and Wheat, H. (1996). Orientation selectivity of thalamic input to simple cells of cat visual cortex. Nature 380, 249–252. 10.1038/380249a0.

39. Winer, J.A. (2005). Decoding the auditory corticofugal systems. Hear Res 207, 1–9. 10.1016/j.heares.2005.06.007.

40. Ibrahim, B.A., Murphy, C.A., Yudintsev, G., Shinagawa, Y., Banks, M.I., and Llano, D.A. (2021). Corticothalamic gating of population auditory thalamocortical transmission in mouse. eLife 10. 10.7554/eLife.56645.

41. Semedo, J.D., Zandvakili, A., Machens, C.K., Yu, B.M., and Kohn, A. (2019). Cortical areas interact through a communication subspace. Neuron 102, 249–259.e4. 10.1016/j.neuron.2019.01.026.

42. Semedo, J.D., Jasper, A.I., Zandvakili, A., Krishna, A., Aschner, A., Machens, C.K., Kohn, A., and Yu, B.M. (2022). Feedforward and feedback interactions between visual cortical areas use different population activity patterns. Nat Commun 13, 1099. 10.1038/s41467-022-28552-w.

## Methods references

43. Mianné, J., Chessum, L., Kumar, S., Aguilar, C., Codner, G., Hutchison, M., Parker, A., Mallon, A.-M., Wells, S., Simon, M.M., et al. (2016). Correction of the auditory phenotype in C57BL/6N mice via CRISPR/Cas9-mediated homology directed repair. Genome Med 8, 16. 10.1186/s13073-016-0273-4.

44. Escabí, M.A., and Read, H.L. (2003). Representation of spectrotemporal sound information in the ascending auditory pathway. Biol Cybern 89, 350–362. 10.1007/s00422-003-0440-8.

45. Gill, P., Zhang, J., Woolley, S.M.N., Fremouw, T., and E. Theunissen, F. (2006). Sound representation methods for spectro-temporal receptive field estimation. J Comput Neurosci 21, 5–20. 10.1007/s10827-006-7059-4.

46. Barnstedt, O., Keating, P., Weissenberger, Y., King, A.J., and Dahmen, J.C. (2015). Functional microarchitecture of the mouse dorsal inferior colliculus revealed through in vivo two-photon calcium imaging. J Neurosci 35, 10927–10939. 10.1523/JNEUROSCI.0103-15.2015.

47. Lu, E., Llano, D.A., and Sherman, S.M. (2009). Different distributions of calbindin and calretinin immunostaining across the medial and dorsal divisions of the mouse medial geniculate body. Hear Res 257, 16–23. 10.1016/j.heares.2009.07.009.

48. Sahani, M., and Linden, J.F. (2003). How linear are auditory cortical responses? Adv. Neural Inf. Process. Syst., 109–116.

49. Park, I.M., Meister, M.L.R., Huk, A.C., and Pillow, J.W. (2014). Encoding and decoding in parietal cortex during sensorimotor decision-making. Nat Neurosci 17, 1395–1403. 10.1038/nn.3800.

