## Supplementary Figures for "Subcortical origin of nonlinear sound encoding in auditory cortex"

### Supplementary Information for Lohse et al., 2024

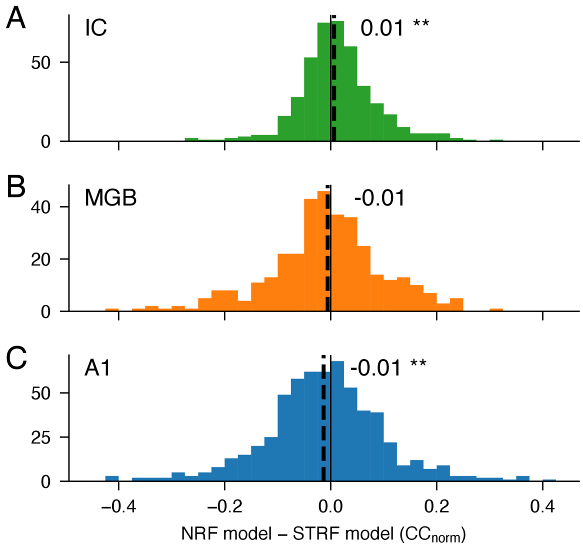


**Figure S1: Neural network models perform similarly to linear-nonlinear STRF models on this dataset**

(A-C) Relative performance ($CC_{norm}$) of Network Receptive Field (NRF) models and linear-nonlinear STRF models in predicting responses of neurons in IC, MGB, A1 respectively. In this study, we used linear-nonlinear STRF models as our baseline for evaluating the performance of population communication models. It is conceivable, therefore, that the apparently high predictive power of population communication models merely reflects poor performance of our STRF models. To ensure this is not the case, we also fitted NRF models, which describe the responses of each neuron using a neural network whose input is the time-varying cochleagram (log-spectrogram). We find that, for this dataset (where the stimuli are spectrally random), the NRF model outperforms the STRF model for some neurons, and underperforms for others, but there are only minor differences in predictive power across the neural population in all three areas.

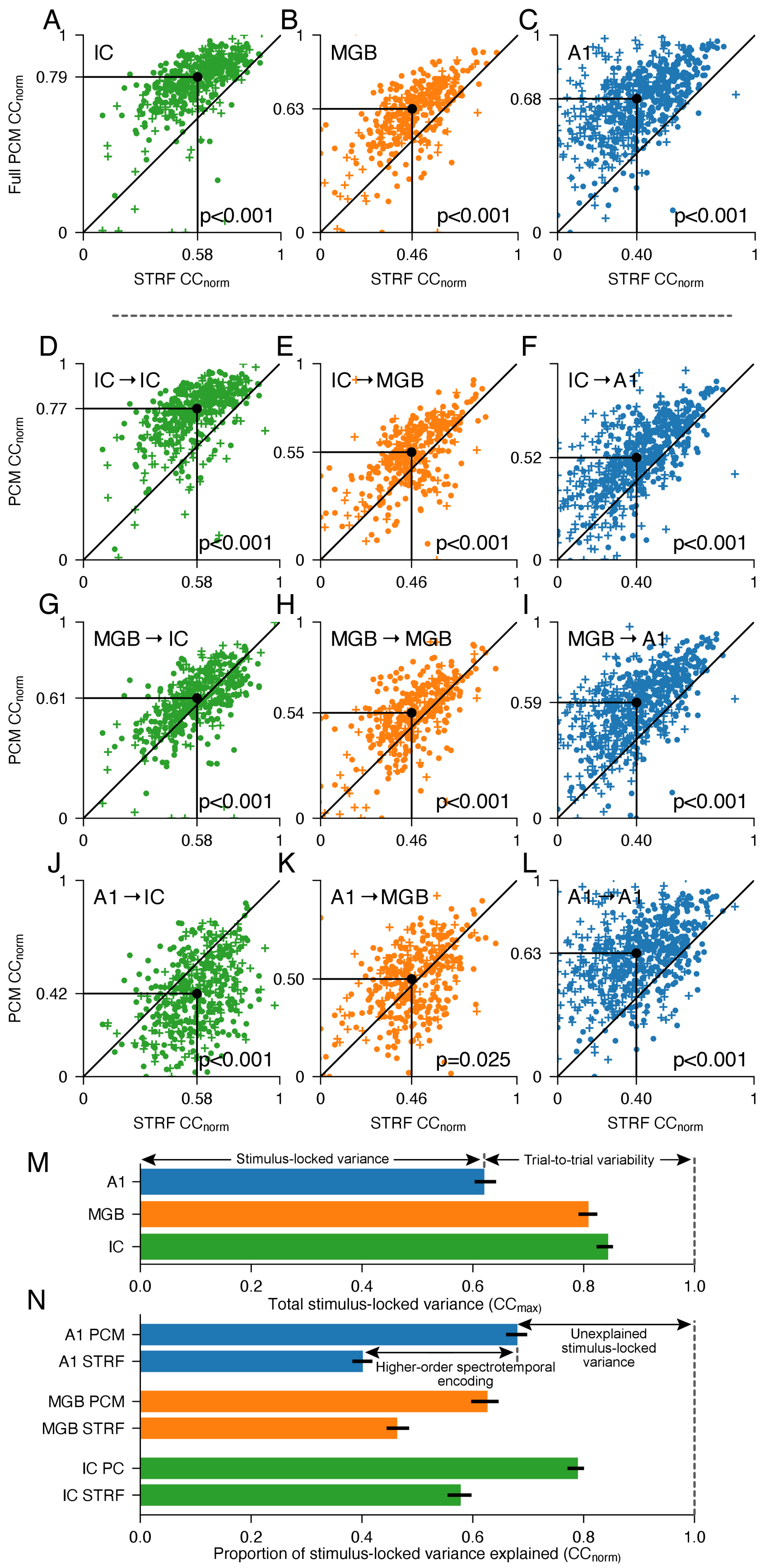


**Figure S2: Additional variance explained by the population communication models, compared to STRF models**

(A-C) Comparison of prediction performance of full population communication model (taking inputs from non-simultaneously-recorded units in IC, MGB and A1; y-axis), relative to STRF model (x-axis), for predicting responses in IC, MGB and A1 respectively.
(D-L) Comparison of prediction performance of single-area population communication models (y-axis), relative to STRF model (x-axis). The source population and target areas are shown in the top-left of each plot (source population → target).
(M) Total stimulus-locked variance ($cc_{max}$) in IC, MGB and A1, respectively. The difference from 1 is a measure of how much variance in each area is not stimulus-locked, i.e., trial-to-trial variability in neural responses.
(N) Median proportion, $cc_{norm}$, of the stimulus-locked variance ($cc_{max}$) that can be explained by STRF and population communication models for each brain region. The performance of the STRF model is a measure of how much variance can be explained by simple spectrotemporal models. The additional performance of the population communication model indicates the proportion of neuronal responses that can be explained by higher-order spectrotemporal encoding, which is stimulus-locked but cannot be modeled with simple spectrotemporal models. The remaining variance is stimulus-locked, but remains unexplained.

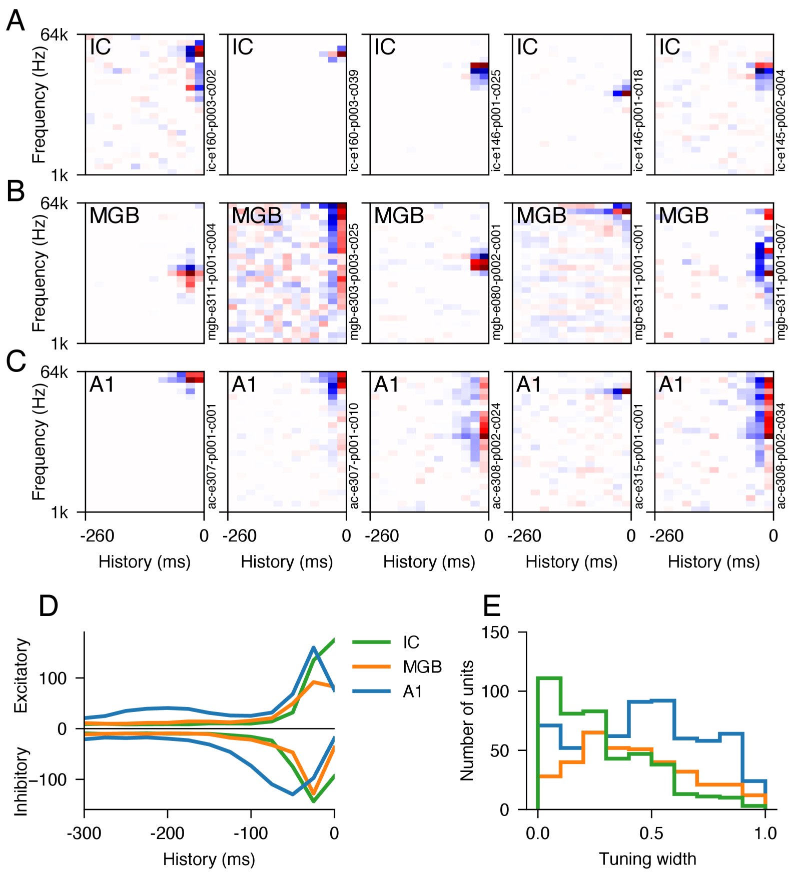


**Figure S3: STRF model characteristics over the auditory hierarchy**
(A-C) Examples of STRFs in IC, MGB and A1, respectively.
(D) Time course of excitatory (top) and inhibitory (bottom) coefficients, summed across all STRFs for each brain area, showing the increase in response latency at higher levels of the auditory hierarchy.
(E) Tuning width of STRFs (see Methods), showing an increase across the hierarchy.

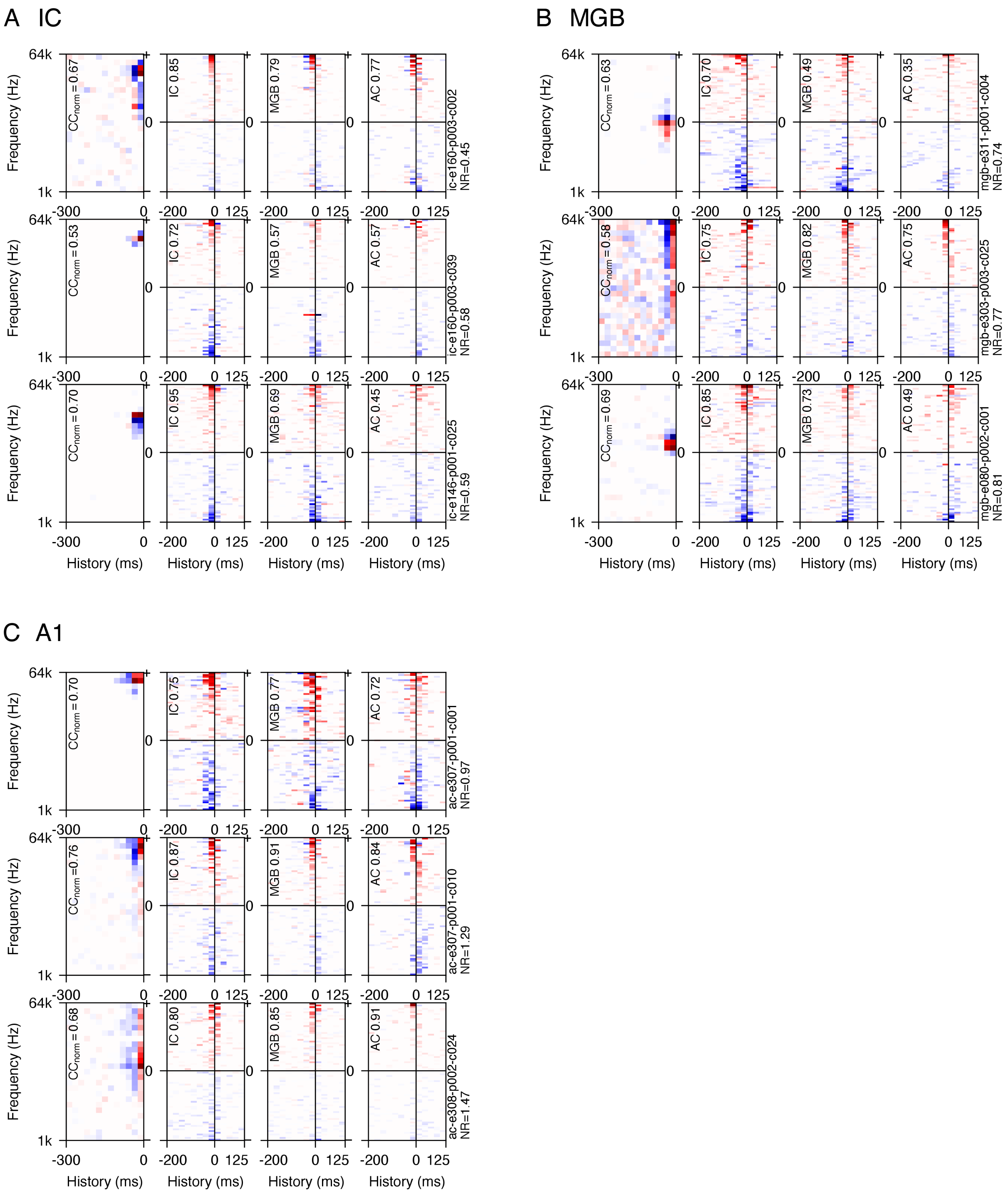


**Figure S4: Comparison of STRF and population communication model kernels over the auditory hierarchy**

(A) Examples of STRFs and population communication model kernels for three example IC neurons. The left column shows the STRF model kernel, and the subsequent three columns show population communication model kernels using IC, MGB and A1 units as regressors (source population), respectively. $CC_{norm}$ values for each model are shown in the top left. For population communication model kernels, only the regressor units with the 20 highest (above the axis) and 20 lowest (below the axis) summed coefficient values are shown, in descending order of summed coefficient value.
(B-C) Similar examples for MGB and A1 units, respectively.

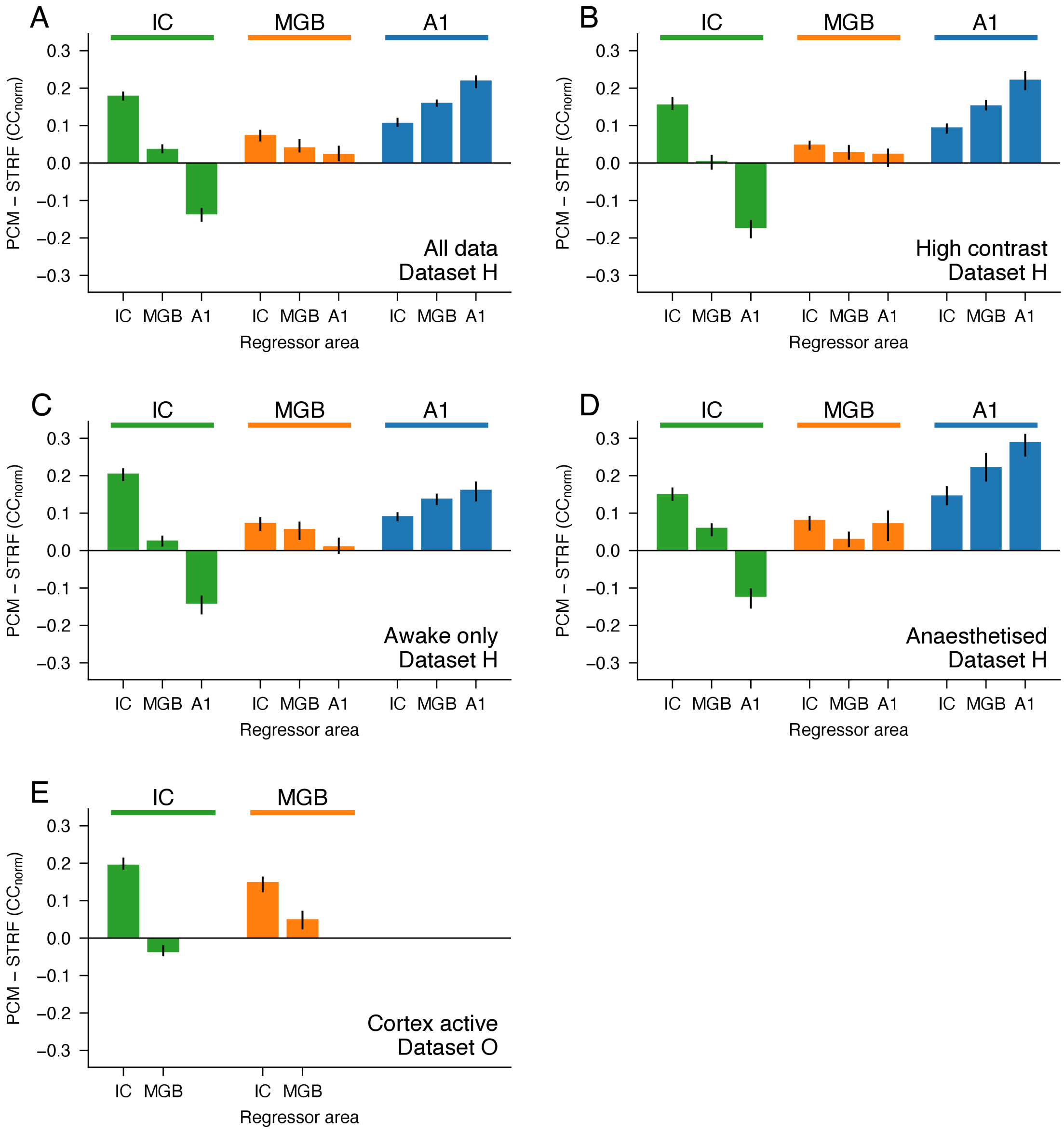


**Figure S5: Control analyses on subsets of data**

Difference (median) between prediction accuracy of population communication models and STRF models for units recorded in the IC, MGB and A1 (target population) with regressor inputs (source population) from different processing levels. The values in these panels are comparable to those in **Figure 3D**, but are measured for different subsets of the data.

(A) Main results for dataset H, as in **Figure 3D**.

(B) Equivalent analysis for dataset H, but using only neural responses to high contrast sounds.

(C) Equivalent analysis including only awake data from dataset H.

(D) Equivalent analysis including only anesthetised data from dataset H.

(E) Equivalent analysis using dataset O (optogenetic data), including only the condition where cortex was active.

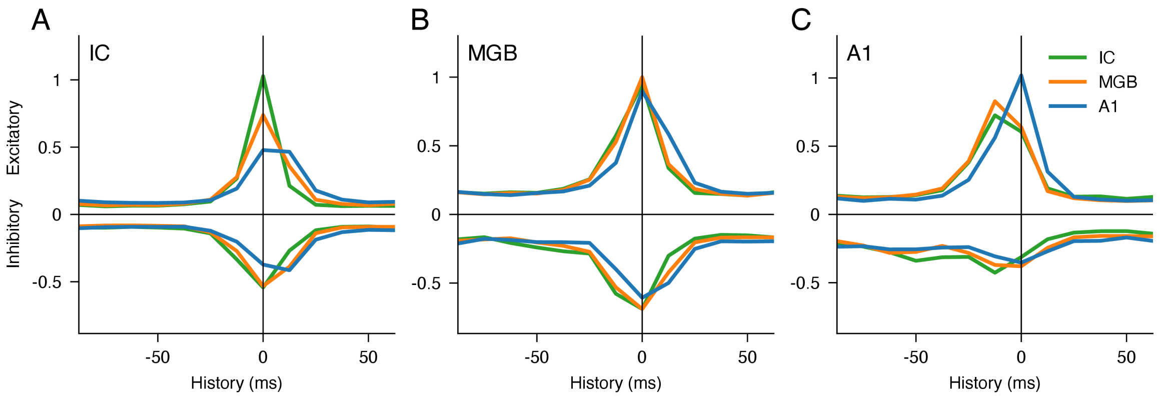


**Figure S6: Time course of population communication models over the auditory processing hierarchy**
(A) Time course of excitatory (above x-axis) and inhibitory (below x-axis) coefficient values in population communication models of IC neurons, summed over regressor units (source population) in single-area population communication models. The mode offset for IC-to-IC coefficient values is 0, and increases for regressor units located in higher auditory areas (MGB and A1). This is expected, given the increasing latency of typical neural responses across the auditory pathway.
(B-C) Similar plots for MGB and A1 target units, respectively.

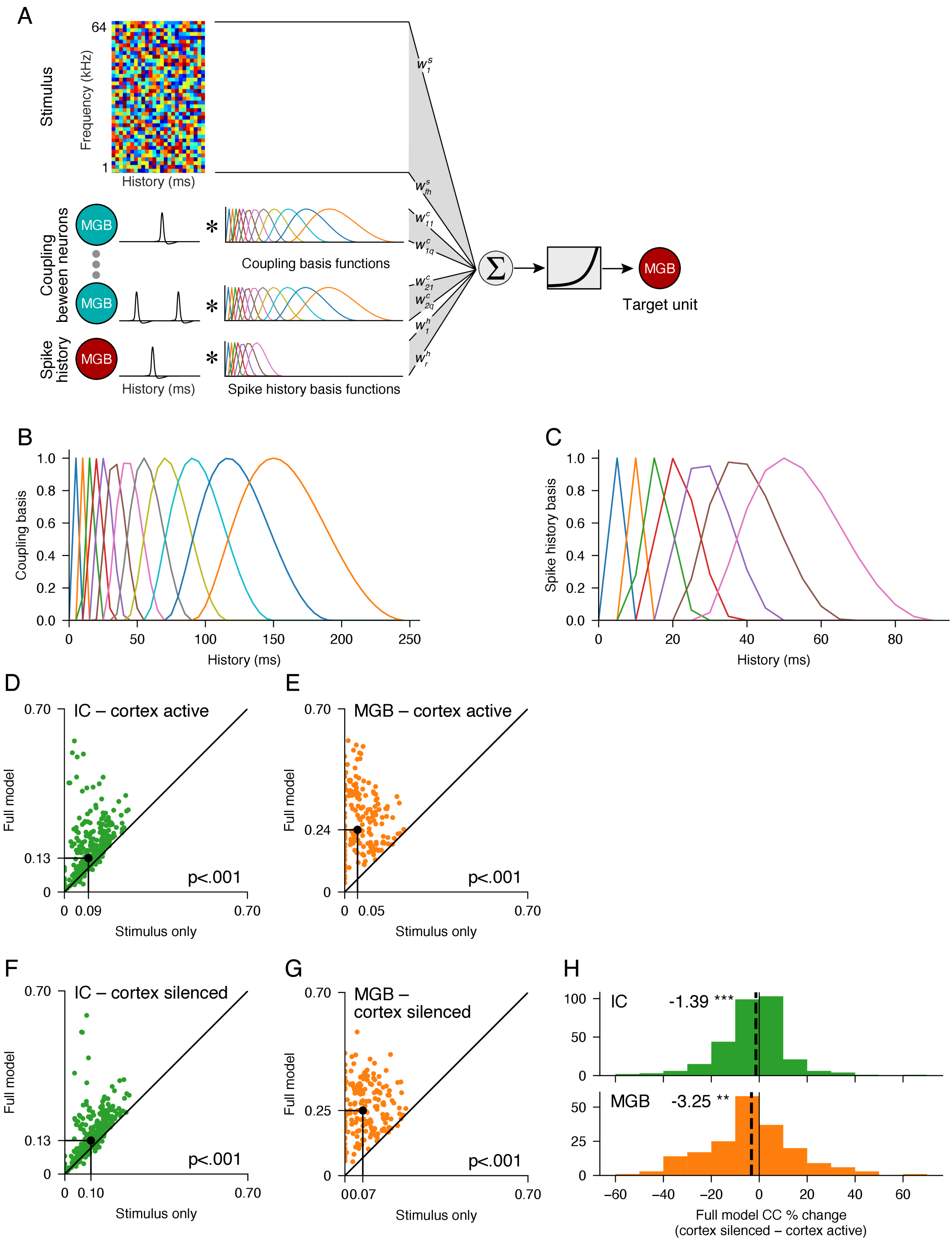


**Figure S7: GLM analysis of neuron-to-neuron communication**

(A) Schematic showing the full GLM model, which consists of an STRF component (similar to STRF models), with additional regressors describing coupling between simultaneously-recorded neurons and the spike history of the target neuron. The spike coupling regressors are the result of convolving the responses of simultaneously-recorded neurons with basis functions (B) spanning latencies up to 250 ms at 5 ms resolution. The spike history regressors are the result of convolving the target

neuron's responses with basis functions (C) spanning latencies up to 90ms. To fit the neural responses, weights are estimated for all regressors which minimise the MSE between the model output and the target neuron's responses, using a Poisson GLM[31].
(D-G) The full model, including coupling and spike history, predicts neuronal responses better than the same model including the stimulus only, in both IC (D, F) and MGB (E, G), and both when the cortex was active (D, E) and when it was silenced optogenetically (F, G).
(H) Performance of the full model is slightly higher when the cortex was active than when the cortex was silenced, for both IC and MGB.
